# The Rab GEF VINE couples phosphatase recruitment to GAP-mediated Rab5 inactivation

**DOI:** 10.64898/2026.01.27.702132

**Authors:** Mia S. Frier, Shawn P. Shortill, Michael Davey, Elizabeth Conibear

## Abstract

Rab5-family GTPases cycle between active and inactive forms to regulate endosomal membrane identity and protein trafficking. VPS9-family guanine nucleotide exchange factors (GEFs) promote Rab activation at endosomes, whereas GTPase-activating proteins (GAPs) oppose Rab signaling. Here, we identify an unexpected role for the yeast VPS9-family GEF complex VINE in promoting the inactivation of the Rab5 homolog Vps21. Through genome-wide proximity screening, predictive modeling, targeted mutagenesis and *in vivo* assays, we show that VINE recruits the protein phosphatase Glc7 through the ankyrin repeat-containing domain of its GEF subunit Vrl1. Our results suggest this directs the dephosphorylation of Kxd1, a subunit of the GAP adaptor BLOC-1, which in turn enhances its interaction with the Vps21-specific GAP Msb3 and accelerates GAP-mediated Vps21 inactivation. Thus, VINE is a VPS9-family GEF complex that selectively limits endosomal Rab signaling. These findings reveal a novel mechanism integrating positive and negative Rab regulation, providing insight into how Rab5 signaling is fine-tuned during endosomal trafficking and maturation.

## Introduction

Protein sorting at endosomes maintains the correct distribution of membrane proteins within the endolysosomal system, enabling proteostasis and signaling essential for cellular health (Yarwood et al., 2020). Protein trafficking from endosomes depends on Rab GTPases, including Rab5 and closely related family members, which control endosomal biogenesis, identity, and maturation (Barr, 2013).

The cycling of Rab5-family GTPases between active and inactive states is controlled by guanine nucleotide exchange factors (GEFs), which promote GTP loading, and GTPase-activating proteins (GAPs), which stimulate GTP hydrolysis. These regulators can display selectivity toward specific Rab isoforms and associate with membranes through interactions with specific lipids, cargo, or components of the trafficking machinery (Delprato et al., 2004; Zhang et al., 2006; Lachmann et al., 2012; Nakashima et al., 2023; Frasa et al., 2012; Lamb et al., 2015; Mattera and Bonifacino, 2008; Kälin et al., 2015; Lauer et al., 2019). In their active GTP-bound form, Rabs recruit effectors that promote vesicle tethering and fusion and enable cargo sorting into anterograde and retrograde trafficking pathways (Murray et al., 2016; Tremel et al., 2021). Endosomal maturation is followed by cargo transfer to the lysosome or vacuole. This is facilitated by a Rab cascade, in which GAP-mediated Rab inactivation is coordinated with the GEF-mediated activation of a downstream-acting Rab GTPase (Rink et al., 2005; Suda et al., 2013; Hiragi et al., 2022; Wilmes and Kummel, 2023).While mechanisms of Rab activation and inactivation have been described in detail, how these opposing activities are coordinated on endosomal membranes to precisely control Rab signaling remains an important unresolved question.

Rab5 proteins are activated by the VPS9 family of Rab GEFs, which is conserved across eukaryotes. Most organisms express multiple VPS9-family proteins that catalyze nucleotide exchange on Rab5 and closely related GTPases (Carney et al., 2006; Herman et al., 2018). Although VPS9-family GEFs share a conserved catalytic domain, they differ in domain architecture, interaction partners, and Rab isoform preferences, suggesting that individual VPS9-family members regulate distinct trafficking events (Kajiho et al., 2003; Sato et al., 2005; Zhang et al., 2014; Shideler et al., 2015; Hsu et al., 2018).

Consistent with this idea, both anterograde and retrograde trafficking pathways have been associated with specific VPS9-family GEFs. In anterograde trafficking, human Rabex5 and its yeast homolog Vps9 interact with ubiquitylated proteins to couple Rab5 activation with cargo internalization and multivesicular body formation (Mattera and Bonifacino, 2008; Aikawa et al., 2012; Russell et al., 2012; Shideler et al., 2015). In the retrograde direction, VPS9 family members bind retromer, an oligomerizing coat complex that mediates endosomal protein recycling (Hesketh et al., 2014; Bean et al., 2015). For example, the human VPS9-family GEF VARP binds retromer and activates Rab21 to link Rab21 signaling to endosome-to-cell surface recycling (Hesketh et al., 2014). More broadly, retromer interacts with multiple Rab GEFs and GAPs, indicating that retrograde trafficking is coordinated by complex Rab signaling networks (Jia et al., 2016; Antón-Plágaro et al., 2025). These examples illustrate how Rab activation by VPS9-family GEFs is directed by the endosomal protein sorting and trafficking machinery. However, the mechanism by which each VPS9-family GEF helps to spatially and temporally control Rab signaling at endosomes remains poorly understood.

Studies in *Saccharomyces cerevisiae* have provided a powerful framework for understanding the shared and distinct contributions of GEFs to endosomal functions. Yeast express three VPS9-family GEFs which together regulate the three Rab5-family GTPases Vps21, Ypt52 and Ypt53. Two of these GEFs, Vps9 and Muk1, exhibit similar nucleotide exchange activity toward each Rab5 isoform *in vitro* (Paulsel et al., 2013; Cabrera and Ungermann, 2013), yet they exert distinct effects on endosomal processes *in vivo*. While multivesicular body formation depends largely on Vps9 (Shideler et al., 2015), autophagy is primarily driven by Vps9 and the major Rab5 homolog Vps21, with Muk1 and other Rab5-family members playing more minor roles (Chen et al., 2014). These findings indicate that specific Rab-GEF pairings can confer distinct endosomal functions.

The third yeast VPS9-family GEF, Vrl1, a homolog of human VARP, has been less well characterized. We recently identified Vrl1 as a subunit of the endosomal VINE complex (Shortill et al., 2022), but whether it preferentially regulates a specific Rab5 isoform is not known. This gap is due in part to a mutation in the *VRL1* gene that prevents expression of full-length Vrl1 in most laboratory strains of *S. cerevisiae* (Bean et al., 2015; Shortill et al., 2022). Because yeast lack direct homologs of the VARP substrate Rab21 and VARP binding partners Rab32 and Rab38, and Vrl1 lacks the Zn-fingernail domain used by VARP to engage retromer, the functions of VINE are unlikely to be directly equivalent to those of VARP (Zhang et al., 2006; Ohbayashi et al., 2012; Tamura et al., 2009; Crawley-Snowdon et al., 2020). Nevertheless, key features are conserved: the N-terminal VPS9 domain and ankyrin repeat-containing regions of VARP are present in Vrl1, and VINE contains a paralog of the retromer membrane adaptor Vps5 (Shortill et al., 2022; Shortill et al., 2024). These similarities raise the possibility that VINE functions as a specialized Rab regulatory platform analogous to the VARP-retromer assembly.

Here, we show that the GEF complex VINE promotes the activity of a Rab GAP, thereby coupling positive and negative regulation of endosomal Rab GTPases. We demonstrate that VINE recruits a phosphatase that enhances GAP-mediated inactivation of Vps21, but not other Rab5-family GTPases, through its action on the GAP adaptor BLOC-1. Together, our findings reveal that VINE fine-tunes endosomal Rab signaling by acting both as a Rab GEF and as a coordinator of Rab inactivation during endosomal maturation.

## Results

### A VINE-specific role in Rab5 regulation

We hypothesized that if VINE makes unique, isoform-selective contributions to endosomal Rab GTPase regulation, it might differentially alter the subcellular localization of the yeast Rab5 isoforms. When overexpressed in wild type laboratory yeast that lack VINE due to a frameshift in the *VRL1* gene (Bean et al., 2015; Shortill et al., 2022), mCherry-tagged Rab5-family proteins Vps21, Ypt53 and Ypt52 localized not to typical small, isolated endosomes, but instead to bright puncta (Gerrard et al., 2000) (Figure 1A). These structures could represent clusters of enlarged endosomes, a hallmark of Rab5 hyperactivation (Russell et al., 2012). Strikingly, when we introduced a functional copy of the *VRL1* gene to allow formation of the VINE complex, overexpressed mCherry-Vps21 no longer appeared primarily at enlarged endosomal structures but instead localized to small, dispersed puncta (Figure 1A). This resulted in a decrease in the number of bright mCherry-Vps21 puncta per cell detected by automated image quantitation (Figure 1B). *VRL1* expression induced a small amount of mCherry-Vps21 to localize in a pattern resembling the nuclear ER (Figure S1A), consistent with increased levels of inactive, GDP-bound Vps21 (Cabrera and Ungermann, 2013). Notably, VINE also reduced the number of puncta brightly labelled by sfGFP-Vps21 expressed under its native promoter from its endogenous genomic locus, suggesting VINE regulates Vps21 even when Vps21 is present at endogenous levels (Figure S1B, C). We observed a similar trend for the Vps21 paralog Ypt53; although posttranscriptional downregulation of *YPT53* limits its overexpression (Schmidt et al., 2017), a small number of enlarged endosomal structures formed in response to Ypt53 overexpression, and VINE reduced this (Figure 1A, B). Notably, the enlarged endosomes formed by overexpression of the other Rab5-family protein, Ypt52, were unaffected by *VRL1* expression, suggesting VINE may promote the inactivation of Vps21 but not Ypt52 (Figure 1A, B).

**Figure 1.**
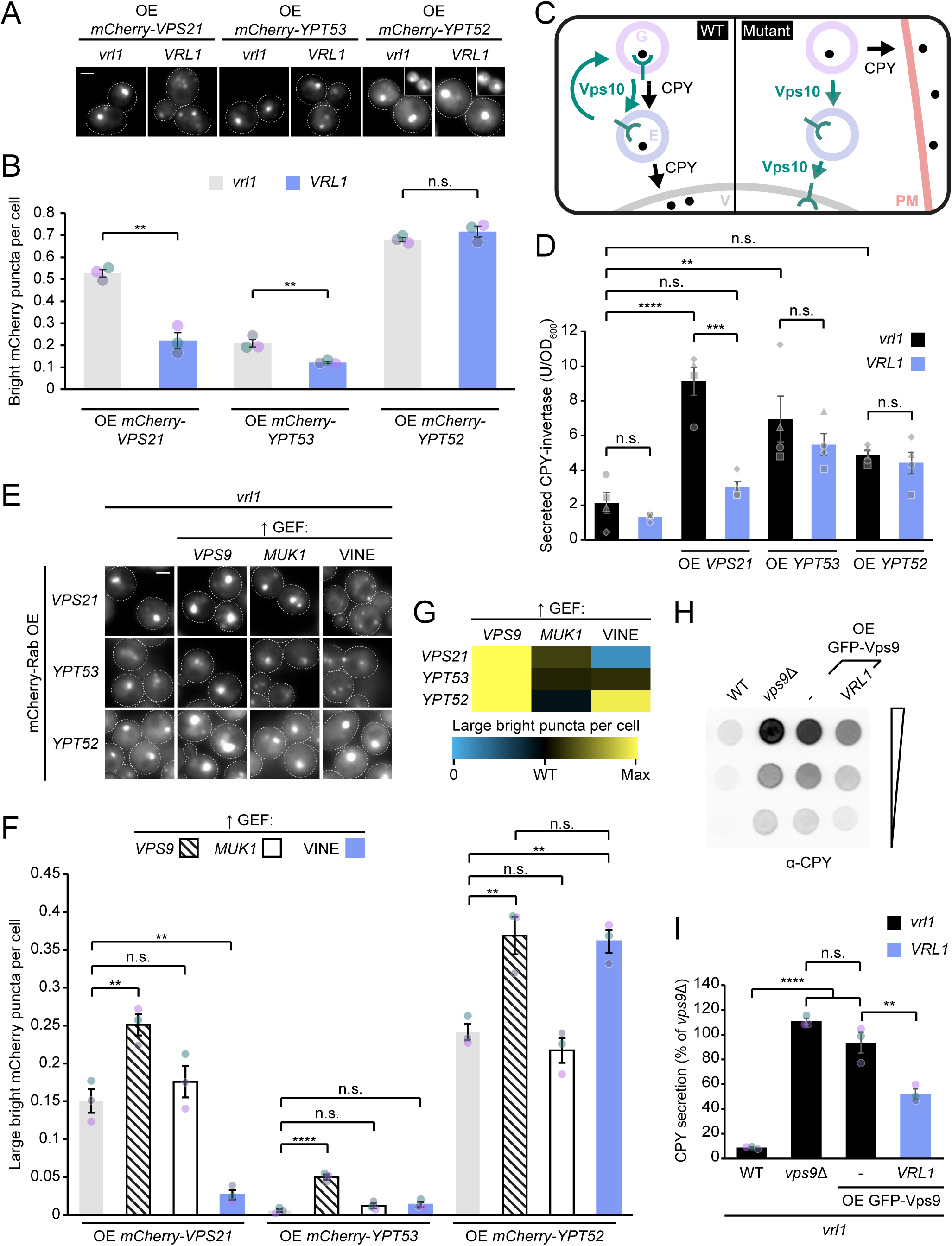
VINE rescues endosomal defects induced by excess active Vps21. (A) VINE selectively rescues the abnormal endosomal morphology caused by overexpression of yeast Rab5 homologs. Fluorescence micrographs of mCherry-Rab fusions overexpressed from the strong *TEF2* promoter in strains lacking Vrl1 (*vrl1*) or with Vrl1 expressed from a plasmid under the *VRL1* promoter. (B) Automated quantitation of bright mCherry puncta per cell, shown in *A*. Unpaired two-tailed equal variance t tests; n=3, cells/strain/replicate ≥ 1231; n.s. = p > 0.05, * = p < 0.05, ** = p < 0.01. (C) Schematic of trafficking pathway mediating delivery of CPY to vacuoles, and CPY mis-trafficking upon endosomal dysfunction. Letters denote organelles: G, Golgi; E, endosome; V, vacuole; PM, plasma membrane. (D) VINE rescues vacuolar cargo missorting caused by *VPS21* overexpression. Quantitation of extracellular activity of secreted invertase-CPY fusion protein. One-way ANOVA with Tukey’s multiple comparisons test; n=4; n.s. = p > 0.05, ** = p < 0.01, *** = p < 0.001, **** = p < 0.0001. (E) The three VPS9-family GEFs have distinct effects on Rab5-family GTPases. Fluorescence micrographs of cells expressing mCherry-tagged Vps21, Ypt53 or Ypt52 from the strong *TEF2* promoter and harboring plasmids expressing HA-tagged Vps9, Muk1 or Vrl1 expressed from the strong *ADH1* promoter. (F) Automated quantitation of large bright puncta labelled by mCherry-Rab fusions in GEF-overexpressing strains, shown in *E*. One-way ANOVA with Tukey’s multiple comparisons test; n=3, cells/strain/replicate ≥ 1131; n.s. = p > 0.05, ** = p < 0.01, **** = p < 0.0001. (G) Heatmap summarizing effects of GEF overexpression on localization of each mCherry-Rab fusion to large bright puncta. (H) VINE partially rescues vacuolar cargo missorting caused by *VPS9* overexpression. Western blot detection of CPY secreted from cells due to *VPS9* deletion or overexpression. (I) Quantitation of secreted CPY in *H*. One-way ANOVA with Tukey’s multiple comparisons test; n=3; n.s. = p > 0.05, ** = p < 0.01, **** = p < 0.0001. Scale bars, 2 µm. Error bars report SEM. OE, overexpressed. CPY, carboxypeptidase Y. U, units of activity. OD_600_, optical density at 600 nm. GEF, guanine nucleotide exchange factor.

If these morphological defects reflect impaired endosomal function, they should be accompanied by defects in cargo sorting. Retromer-mediated retrograde trafficking of the carboxypeptidase Y (CPY) receptor Vps10 from endosomes maintains pools of Vps10 both at endosomes and the Golgi (Cooper and Stevens, 1996). We found that overexpressing mCherry-tagged Rab5 isoforms caused GFP-tagged Vps10 to colocalize with Rab5 proteins at enlarged endosomal structures (Figure S1D). Localization of Vps10 exclusively to endosomal structures is consistent with delayed retrograde trafficking from endosomes, which could impair delivery of the Vps10 ligand CPY from the Golgi to vacuoles (Figure 1C). Indeed, overexpressing untagged Vps21 caused a significant increase in the secretion of a CPY-invertase fusion protein, consistent with a mild defect in Vps10 endosome-to-Golgi trafficking, which was rescued by *VRL1* expression (Figure 1D). This supports a model in which VINE promotes Vps21 inactivation, an unexpected activity for a VPS9-family GEF.

To determine if this activity is unique to VINE, we tested whether overexpression of other VPS9-family GEFs affected the localization of overexpressed mCherry-tagged Vps21, Ypt53 and Ypt52. Vps9 overexpression increased the number of bright mCherry-Rab puncta regardless of which Rab5-family protein was overexpressed (Figure 1E-G). Vps9 GEF activity, but not the Vps9 CUE domain, was required for the Vps21-driven formation of enlarged endosomal structures (Figure S1E, F). Muk1 overexpression did not influence this phenotype (Figure 1E-G) and even when expressed as a fusion with the Vps9 CUE domain, which can target Muk1 to endosomes (Shideler et al., 2015), was unable to support Vps21-driven enlarged endosome formation (Figure S1G, H). Overexpressing GFP-Vps9 also caused CPY secretion, indicating defects in endosomal protein sorting (Figure 1H, I). *VRL1* expression partially rescued this, indicating VINE can rescue endosomal protein sorting defects driven by increased endosomal Rab activation. Together these results indicate that the Vps21-inactivating function of VINE is not shared by other VPS9-family GEFs, and that while excess Vps9-mediated Rab activation can drive endosomal defects, VINE can oppose this.

### VINE negatively regulates Vps21 independently of the Vrl1 GEF active site

That VINE can negatively regulate Vps21 was surprising, because Vrl1 is a member of the VPS9 family which promotes the activity of Rab5-family GTPases (Bean et al., 2015; Shortill et al., 2022). We therefore sought to determine the mechanism by which VINE suppresses Vps21 function. First, we asked whether increased Ypt52 activation by VINE might impinge on Vps21 signaling, forming the basis for the observed suppression of Vps21. However, VINE was still able to decrease the localization of mCherry-Vps21 to enlarged endosomes despite deletion of the *YPT52* gene, suggesting VINE does not leverage a Ypt52-dependent pathway to oppose Vps21 function (Figure S2A, B). We next wondered whether the GEF activity of Vrl1 is required for Vps21 inactivation. Expressing an allele with a mutation in the catalytic site of the VPS9 domain (Vrl1^D373A^) decreased the number of bright Vps21 puncta, revealing a GEF activity-independent role of VINE in Rab regulation (Figure 2A, B).

**Figure 2.**
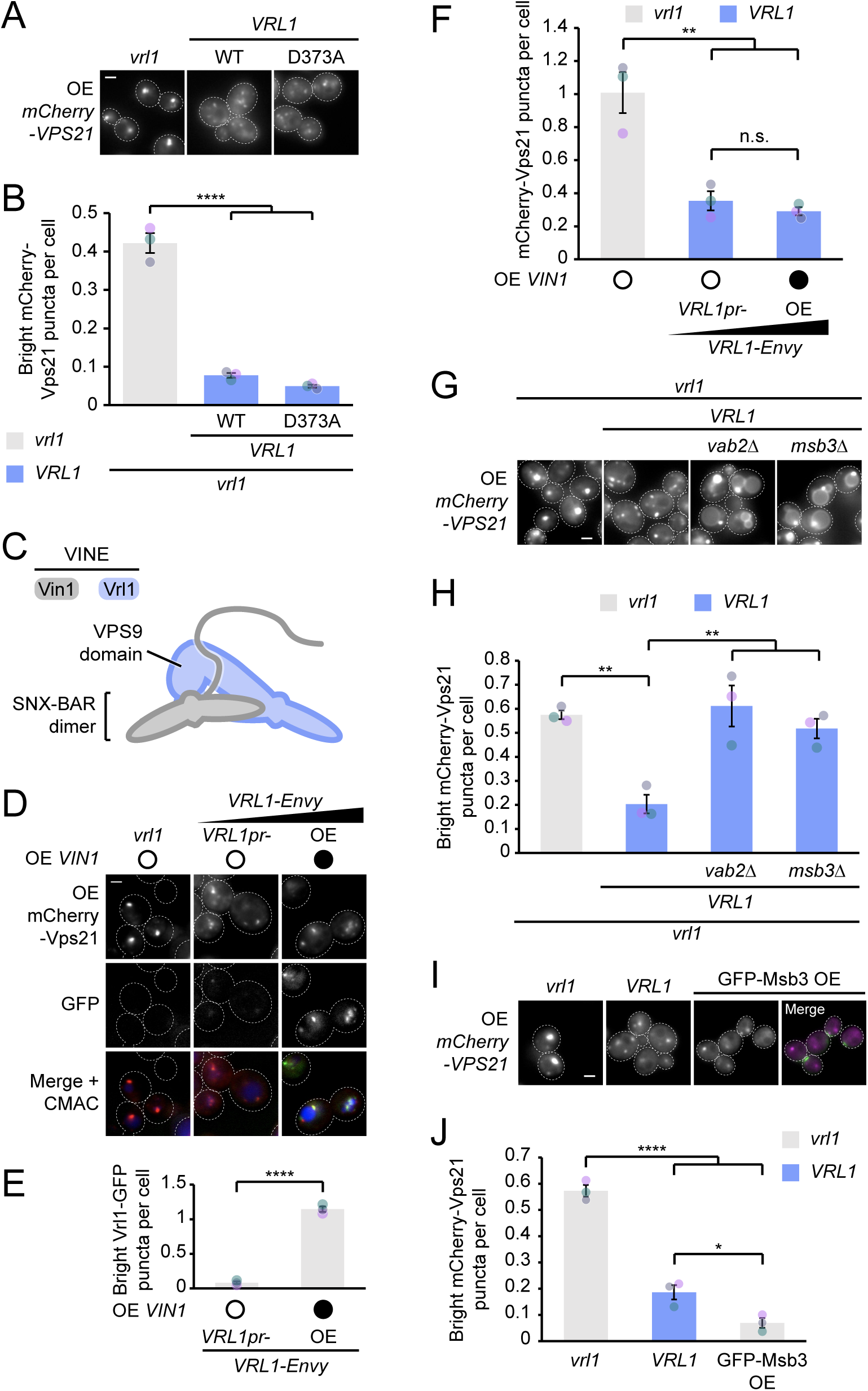
VINE activates the Vps21 GAP independently of the Vrl1 GEF active site. (A) Vrl1 does not require its GEF activity to negatively regulate Vps21. Fluorescence micrographs of cells co-expressing mCherry-Vps21 under control of the *TEF2* promoter, and Envy-tagged WT or GEF-inactive Vrl1 alleles from plasmids under the *VRL1* promoter. (B) Automated quantitation of bright mCherry-Vps21 puncta shown in *A*. One-way ANOVA with Tukey’s multiple comparisons test; n=3, cells/strain/replicate ≥ 1258; **** = p < 0.0001. (C) Schematic of the VINE complex indicating the position of the Vrl1 VPS9 GEF domain relative to the SNX-BAR domains of Vrl1 and Vin1. (D) The endogenous level of VINE is sufficient for Vps21 suppression. Fluorescence micrographs of CMAC-stained live cells co-expressing mCherry-Vps21 from the *TEF2* promoter and endogenous levels of VINE (plasmid-borne Vrl1-Envy expressed from the *VRL1* promoter and Vin1 expressed from its endogenous locus) or elevated levels of VINE (Vrl1-Envy expressed from the strong *ADH1* promoter, with an extra copy of *VIN1* overexpressed from the *RPL18B* promoter). (E) Automated quantitation of bright Vrl1-GFP puncta in *D*. Two-tailed equal variance t test; n=3, cells/strain/replicate ≥ 660; **** = p < 0.0001. (F) Automated quantitation of bright mCherry-Vps21 puncta in *D*. One-way ANOVA with Tukey’s multiple comparisons test; n=3, cells/strain/replicate ≥ 1254; n.s. = p > 0.05, ** = p < 0.01. (G) VINE requires Msb3 and its adaptor BLOC-1 to negatively regulate Vps21. Fluorescence micrographs of the indicated WT or mutant strains co-expressing mCherry-Vps21 from the *TEF2* promoter and Vrl1-Envy from the *ADH1* promoter. (H) Automated quantitation of bright mCherry-Vps21 puncta from *G*. One-way ANOVA with Dunnett’s multiple comparisons test; n=3, cells/strain/replicate ≥ 1599; ** = p < 0.01. (I) VINE or excess GAP is sufficient to oppose Vps21. Fluorescence micrographs of cells expressing mCherry-Vps21 from the *TEF2* promoter, with untagged Vrl1 expressed from the *VRL1* promoter or with GFP-Msb3 expressed from its endogenous locus under the strong *NOP1* promoter. Merge shows mCherry-Vps21 in magenta and GFP-Msb3 in green. (J) Automated quantitation of bright mCherry-Vps21 puncta in *I*. One-way ANOVA with Tukey’s multiple comparisons test; n=3, cells/strain/replicate ≥ 929; n.s. = p > 0.05, **** = p < 0.0001. Scale bars, 2 µm. Error bars report SEM. OE, overexpressed. sfGFP, superfolder GFP. GEF, guanine nucleotide exchange factor. CMAC, 7-amino-4-chloromethylcoumarin.

Because Vrl1 and Vin1 are not highly abundant proteins, we hypothesized that if VINE opposes Vps21 function by directly sequestering Vps21 or a Vps21 effector, higher levels of VINE should more effectively inactivate Vps21. Co-overexpressing both Vrl1 and Vin1 to increase the level of VINE (Figure 2C) resulted in increased endosomal localization of Vrl1-Envy (Figure 2D, E) but did not further decrease the number of bright Vps21 puncta when compared to cells expressing Vrl1 and Vin1 from their endogenous promoters (Figure 2D, F). This indicates that a low level of VINE is sufficient to negatively regulate Vps21, suggesting VINE is not simply sequestering excess Vps21 but may instead act through a catalytic mechanism.

Our results indicate that VINE specifically downregulates Vps21 signaling without opposing other Rab5 isoforms. This apparent selectivity mirrors the effects of the GAP Msb3, which is recruited by the BLOC-1 complex to endosomes where it inactivates Vps21 but not Ypt52 (John Peter et al., 2013; Lachmann et al., 2012; Nickerson et al., 2012). Deleting the gene encoding either Msb3 or the BLOC-1 subunit Vab2 caused Vps21 to localize to enlarged endosomes despite *VRL1* expression (Figure 2G, H). VINE also had no obvious effect on the distribution of mCherry-Vps21^Q66L^, which is resistant to inactivation by Msb3 (Figure S2C). This suggests that VINE acts via Msb3 and BLOC-1 to inactivate Vps21. Mirroring the effect of VINE expression, Msb3 overexpression decreased the number of bright mCherry-Vps21 puncta, consistent with a model in which VINE enhances the ability of Msb3 to inactivate Vps21 (Figure 2I, J; John Peter et al. 2013).

### A conserved surface on the Vrl1 ankyrin repeat-containing domain is critical for Vps21 regulation

Because VINE negatively regulated Vps21 independently of Vrl1 GEF activity, we reasoned that another domain in VINE must mediate Vps21 inactivation. Vrl1 possesses an ankyrin repeat-containing domain (AnkRD) that features a binding site for the VINE subunit Vin1 (Shortill et al., 2022). Mapping the conservation of individual amino acid residues onto a partial predicted model of Vrl1 revealed a conserved region on the surface of the AnkRD that was distinct from the Vin1 binding site (Figure 3A-C). Strikingly, mutation of two conserved residues in this region (Asp638 and Asn642, which were mutated to asparagine and serine respectively or to alanine) prevented VINE from redistributing mCherry-Vps21 from enlarged endosomes (Figure 3D, E; Figure S3A) and from rescuing the CPY-invertase secretion caused by Vps21 overexpression (Figure 3F). In contrast, mutating two adjacent lysine residues had no discernible effect on Vps21 regulation by Vrl1 (Figure 3D, E). Vrl1^D638N N642S^ (hereafter referred to as Vrl1^DN-NS^) was able to recruit Vin1-mScarletI to puncta (Figure S3B, C) and co-immunoprecipitated with Vin1-GFP at a level indistinguishable from Vrl1^WT^ (Figure S3D), indicating this conserved AnkRD site is required for negatively regulating Vps21 but not for other functions of Vrl1.

**Figure 3.**
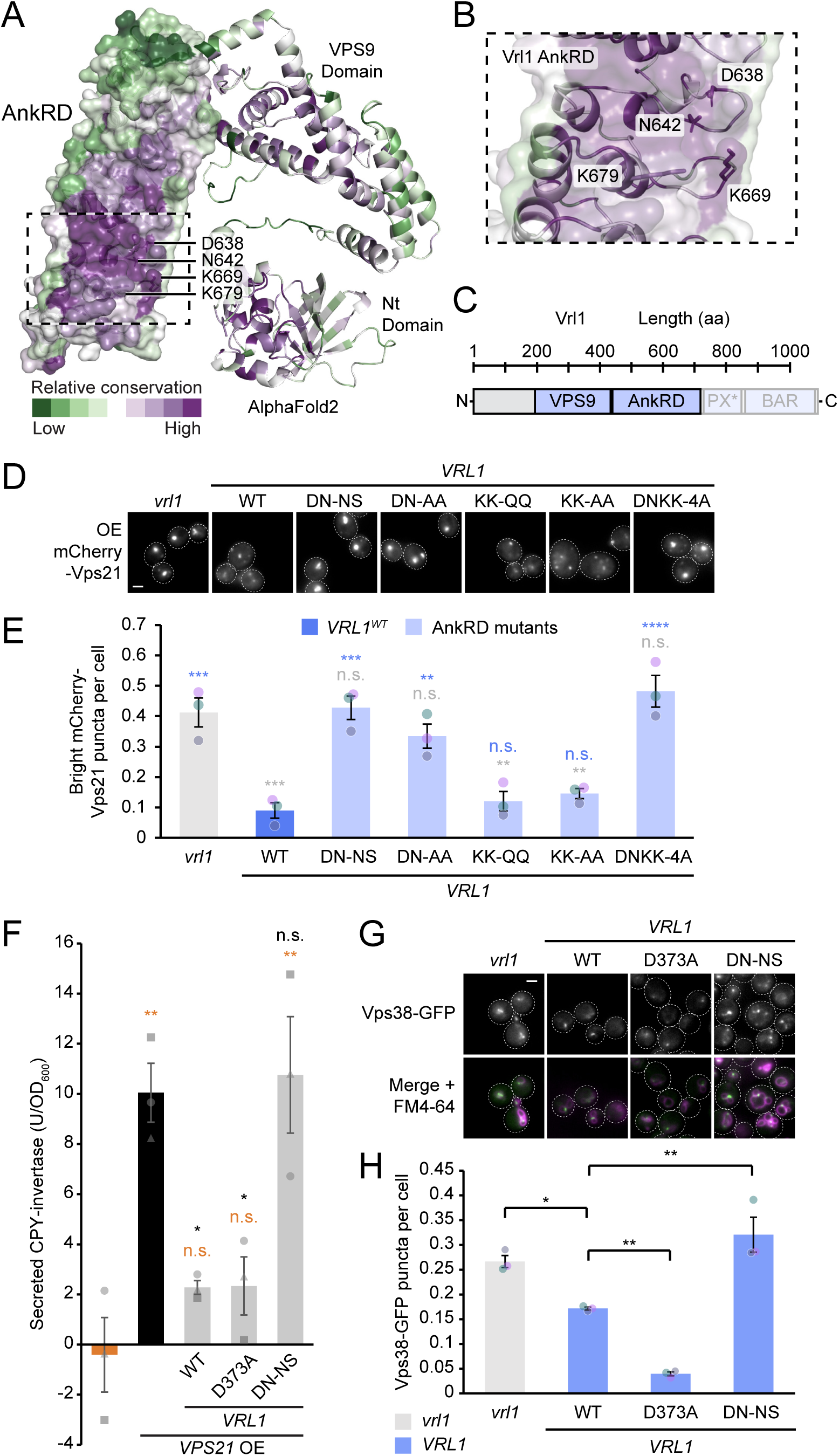
VINE acts via the Vrl1 ankyrin repeat-containing domain to oppose Vps21 function. (A) AlphaFold2-predicted structure of residues 1 to 730 of Vrl1 coloured according to amino acid conservation among fungal homologs using ConSurf (Ashkenazy et al., 2016). Conserved residues mutated in *D* are indicated. Not shown is the Vin1^Nt^-binding site on the opposite face of the AnkRD domain. (B) Detail of *A* showing residues selected for mutagenesis. (C) Schematic of the Vrl1 protein showing the region modeled in *A*. (D) VINE negatively regulates Vps21 via a conserved site on the Vrl1 AnkRD. Fluorescence micrographs of cells expressing *TEF2* promoter-driven mCherry-Vps21 and *ADH1* promoter-expressed Vrl1-Envy harboring mutations in the conserved AnkRD site. (E) Automated quantitation of bright mCherry-Vps21 puncta in E, showing the effects of mutations of the Vrl1 AnkRD. One-way ANOVA with Tukey’s multiple comparisons test; n=3, cells/strain/replicate ≥ 767; n.s. = p > 0.05, ** = p < 0.01, *** = p < 0.001, **** = p < 0.0001. (F) Quantitation of extracellular invertase activity due to secretion of a CPY-invertase fusion in Vps21-overexpressing cells. One-way ANOVA with Tukey’s multiple comparisons test; n=3; n.s. = p > 0.05, * = p < 0.05, ** = p < 0.01, *** = p < 0.001. (G) The Vrl1 GEF active site and the conserved AnkRD site have opposite effects on Vps21 effector localization. Fluorescence micrographs of cells expressing Vps38-GFP from its endogenous locus with or without the expression of untagged Vrl1 mutant alleles. Vacuoles are stained with FM4-64. (H) Automated quantitation of Vps38-GFP puncta in *E*. One-way ANOVA with Dunnett’s multiple comparisons test; n=3, cells/strain/replicate ≥ 1474; * = p < 0.05, ** = p < 0.01. Scale bars, 2 µm. Error bars report SEM. AnkRD, ankyrin repeat-containing domain. Nt, N-terminal. PX*, phox homology-like. BAR, Bin/amphiphysin/Rvs. OE, overexpressed. DN-NS, D638N N642S. DN-AA, D638A N642A. KK-QQ, K669Q K679Q. KK-AA, K669A K679A. DNKK-4A, D638A N642A K669A K679A. CPY, carboxypeptidase Y. U, units of invertase activity. OD_600_, optical density at 600 nm. sfGFP, superfolder GFP.

If the AnkRD site drives Vps21 inactivation, it should oppose the effect of Vrl1 GEF activity on endosomal Rab activity and therefore inhibit effector recruitment. The Rab5 effectors Vps38 and Vac1 can be observed at endosomes when expressed as GFP fusions from their endogenous promoters. A Vrl1 mutant in which the GEF active site is mutated, Vrl1^D373A^, caused a decrease in detectable puncta decorated by the phosphoinositide 3-kinase subunit Vps38-GFP, consistent with Vps21 inactivation through the AnkRD (Figure 3G, H). Conversely, expressing Vrl1^DN-NS^, which has a disrupted AnkRD site, increased Vps38-GFP puncta, consistent with increased Rab activation by the GEF activity of Vrl1. A similar trend was observed for the endosomal tether sfGFP-Vac1 (Figure S3E, F). Together these data suggest that AnkRD-mediated Vps21 inactivation can reduce the pool of active, effector-binding Vps21, while GEF activity can counterbalance this effect.

### The VINE AnkRD associates with Glc7 to drive Vps21 inactivation

To identify how the Vrl1 AnkRD promotes Vps21 inactivation, we sought binding partners specific to this domain. We performed a modified split-dihydrofolate reductase (DHFR)-based genome-wide protein fragment complementation screen to identify proteins that require the Vrl1 AnkRD site for their proximity to VINE. In a standard split-DHFR assay, bait and prey proteins are each tagged with a fragment of the DHFR enzyme in haploid cells, and after mating to form diploids, proximity-driven formation of the functional DHFR enzyme enables growth on media containing methotrexate (Tarassov et al., 2008). In our modified assay, co-expressing a GFP-binding nanobody fused to the C-terminal DHFR fragment (DHFR^C^) allowed us to screen a large library of N-terminal GFP-tagged proteins, expressed at uniform levels from the *NOP1* promoter, for proximity to Vrl1-DHFR^N^ (Fridy et al., 2015; Yofe et al., 2016; Weill et al., 2018) (Figure 4A).

**Figure 4.**
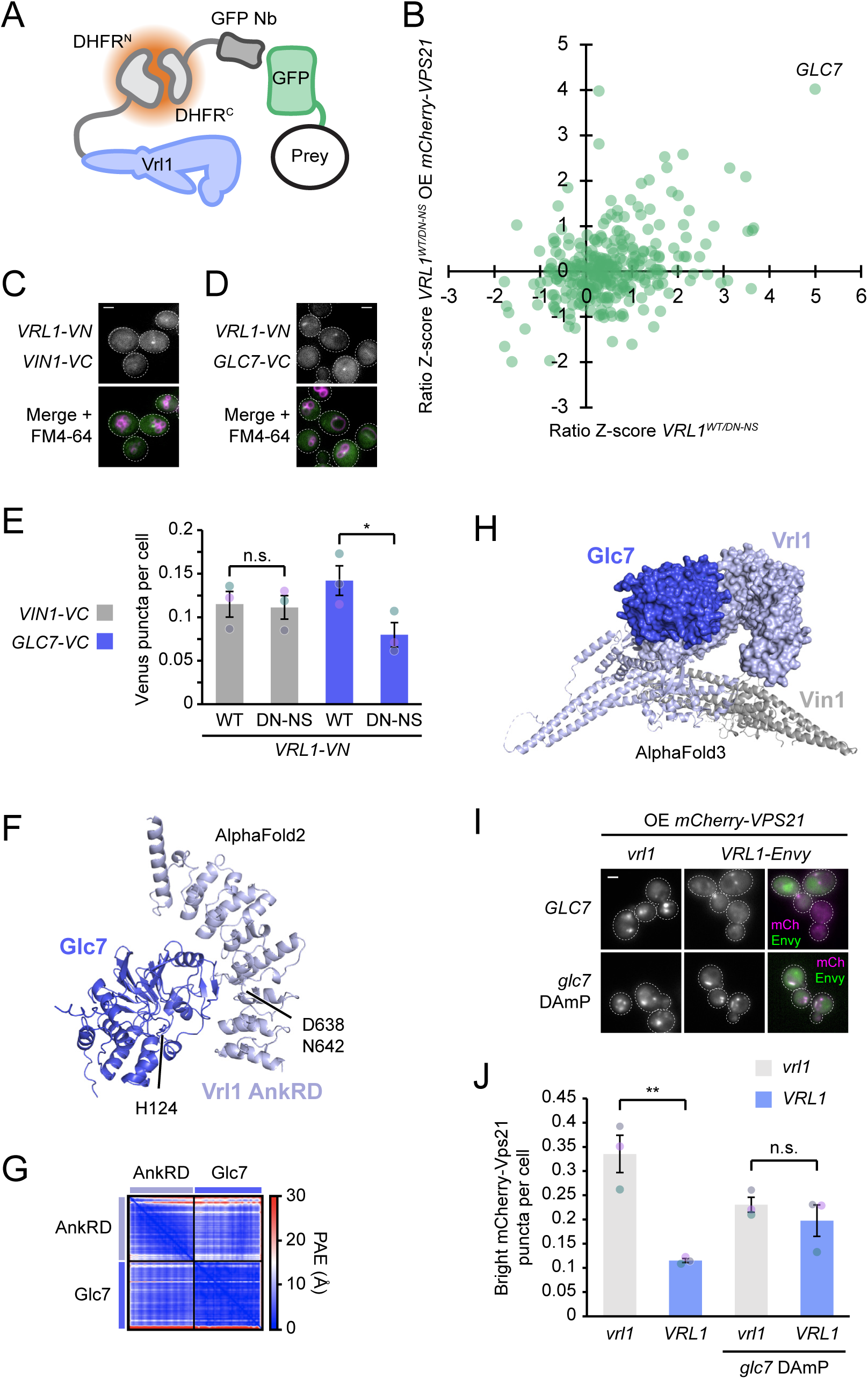
The VINE AnkRD associates with Glc7 to drive Vps21 inactivation. (A) Schematic of modified split-DHFR assay, showing the DHFR^N^-tagged Vrl1 bait and a GFP nanobody (GFP Nb)-DHFR^C^ fusion targeting GFP-tagged prey, wherein bait-prey proximity facilitates reconstitution of a functional DHFR enzyme. (B) The Vrl1 AnkRD specifically mediates proximity to the phosphatase Glc7. Plot of ratio Z-scores from Vrl1 proximity screen. Colony area of strain expressing Vrl1^WT^ was divided by colony area of corresponding strain expressing Vrl1^DN-NS^ to obtain interaction ratio for each prey and converted to Z-scores. The x axis shows the ratio Z-score in strains expressing Vps21 from its endogenous promoter, and the y axis shows the ratio Z-score in strains overexpressing mCherry-Vps21 from the *TEF2* promoter. The AnkRD-dependent hit *GLC7* is indicated. (C) Vrl1 and Vin1 interact at endosomes. Fluorescence micrograph of FM4-64-stained cells expressing Vrl1 and Vin1 as fusions with complementary fragments of Venus. (D) Vrl1 and Glc7 interact at endosomes. Fluorescence micrograph of FM4-64-stained cells expressing Vrl1 and Glc7 as fusions with complementary fragments of Venus. (E) Mutations in the Vrl1 AnkRD selectively disrupt the interaction with Glc7 and not with Vin1. Automated quantitation of Venus puncta per cell in strains shown in *C, D* expressing Vrl1-VN with or without point mutations in the AnkRD and either Vin1-VC (gray; representative image in *C*) or Glc7-VC (indigo; representative image in *D*). Two-tailed equal variance t tests; n=3; cells/strain/replicate ≥ 853; n.s. = p > 0.05, * = p < 0.05. (F) AlphaFold2-predicted interaction of Glc7 with residues 434 to 729 of Vrl1. His 124 of the Glc7 active site and D638 and N642 of the Vrl1 AnkRD are indicated. (G) The predicted interaction between the Vrl1 AnkRD and Glc7 exhibits low error. Plot showing PAE for predicted interaction in *F*. (H) AlphaFold3-predicted interaction of the VINE complex with Glc7. Only residues with pLDDT>50 are shown. (I) VINE requires Glc7 to negatively regulate Vps21. Fluorescence micrographs of *GLC7* or *glc7* DAmP cells co-expressing mCherry-Vps21 from the *TEF2* promoter and Vrl1-Envy. (J) Automated quantitation of bright mCherry-Vps21 puncta in *I*. One-way ANOVA with Tukey’s multiple comparisons test; n=3, cells/strain/replicate ≥ 785; n.s. = p > 0.05, ** = p < 0.01. Scale bars, 2 µm. Error bars report SEM. DHFR, dihydrofolate reductase. SWAT, SWAp-Tag. DN-NS, D638N N642S. OE, overexpressed. VN, Venus N-terminal fragment. VC, Venus C-terminal fragment. AnkRD, ankyrin repeat-containing domain. PAE, predicted aligned error. DAmP, decreased abundance by mRNA perturbation. mCh, mCherry. pTM, predicted template modeling. ipTM, interface predicted TM. Nb, nanobody.

We hypothesized that Vrl1 and a binding partner of the Vrl1 AnkRD site would exhibit a proximity interaction that is impaired by mutations in the AnkRD site. To detect such interactions, we conducted parallel proximity screens using either Vrl1^WT^, or Vrl1^DN-NS^ as bait. We then calculated the ratio of each prey’s DHFR signal with Vrl1^WT^-DHFR^N^ or with the AnkRD mutant Vrl1^DN-NS^-DHFR^N^ and expressed this as a Z-score (see Table S1). In case the AnkRD interactor is recruited upon Rab hyperactivation, we carried out screens both in the presence of endogenous levels of Vps21 and in a strain background expressing mCherry-Vps21 from the strong *TEF2* promoter. One Vrl1 proximity interactor, *GLC7*, gave consistently high Vrl1^WT^/Vrl1^DN-NS^ ratio Z-scores (5.00 when Vps21 was expressed from its endogenous promoter and 4.02 when mCherry-Vps21 was overexpressed), suggesting the AnkRD site is important for proximity to Glc7 (Figure 4B).

Glc7 is the catalytic subunit of protein phosphatase 1 (PP1) in yeast and is recruited to subcellular sites by a suite of adaptor proteins to form PP1 complexes with various essential and nonessential functions (Cannon, 2010). To validate the interaction between VINE and Glc7, we used a split-fluorophore assay that detects proximity interactions between two proteins fused to complementary fragments of Venus (VN and VC). Co-expression of Vrl1-VN with its obligate partner Vin1-VC generated a Venus signal at endosomes proximal to FM4-64-stained vacuoles, as expected (Figure 4C). Consistent with an interaction of VINE and Glc7 at endosomes, co-expressing Vrl1-VN with Glc7-VC also resulted in proximity-induced Venus fluorescence at perivacuolar puncta (Figure 4D). Mutations in the AnkRD impaired the interaction between Vrl1-VN and Glc7-VC but not between Vrl1-VN and Vin1-VC, indicating this site on the Vrl1 AnkRD promotes the interaction between VINE and Glc7 at endosomes but is not required for VINE formation (Figure 4E).

Because our proximity assays suggested the Vrl1 AnkRD mediates an interaction between VINE and Glc7, we sought to visualize how Glc7 might engage the AnkRD. Structural modeling using both AlphaFold2 and AlphaFold3 (Jumper et al., 2021; Abramson et al., 2024) confidently predicted Glc7 in complex with the Vrl1 AnkRD, consistent with Glc7 as a direct binding partner for the Vrl1 AnkRD (Figure 4F, G, Figure S4A). Vrl1 residues Asp638 and Asn642 were found at the predicted interface, consistent with our observations that mutating these residues reduced the proximity of Glc7 to Vrl1. Using *in silico* modeling we also generated a predicted structure of Vrl1, Vin1 and Glc7, with Glc7 situated above the VINE phox homology-Bin/amphiphysin/Rvs (PX-BAR) dimer (Figure 4H). As a PP1 catalytic subunit, Glc7 binds not only to regulatory subunits that direct its localization and promote its activity on specific substrates, but also to inhibitors that prevent substrate recognition or block catalytic activity (Virshup and Shenolikar, 2009). Because the Glc7 active site was predicted to be positioned adjacent to, but not obscured by, the AnkRD (Figure 4F), it is unclear from these models whether VINE might bind to an active or inactive form of Glc7.

To test if Glc7 activity is required to inactivate Vps21, we reduced Glc7 function using a Decreased Abundance by mRNA Perturbation (DAmP) allele that destabilizes the *GLC7* mRNA (Breslow et al., 2008). This allele prevented Vrl1 from decreasing bright mCherry-Vps21 puncta (Figure 4I, J). This suggests that VINE requires Glc7 to negatively regulate Vps21. Together with our data indicating Glc7 interacts with the critical Vps21-suppressing region of the Vrl1 AnkRD, these results support a model in which VINE and Glc7 form an active endosomal PP1 complex whose activity promotes Vps21 inactivation.

Because our data indicate the phosphatase Glc7 acts to ultimately oppose Vps21 activity, we explored whether cognate kinases exist that promote Vps21 function. We considered two kinases as possible candidates: Yck3, which is implicated in Mon1-Ccz1 regulation during the endosomal Vps21-to-Ypt7 cascade (Langemeyer et al., 2020), and the Yck3-regulated kinase Env7 (Manandhar et al., 2020), which was found in our screens to be in close proximity to Vrl1. Although overexpressed Vps21 required neither *YCK3* nor *ENV7* to drive the formation of enlarged endosomal structures (Figure S4B), *ENV7* overexpression did exacerbate this phenotype (Figure S4C, D), suggesting it could be a part of a redundant network of kinases regulating Vps21 activity.

### VINE-dependent Vps21 inactivation requires a secondary interface between Msb3 and BLOC-1

We hypothesized that the novel endosomal PP1 complex formed by VINE and Glc7 acts on proteins known to regulate Vps21. A promising candidate is the GAP Msb3 which has the same substrate selectivity as VINE. Our results (Figure 2G-J) suggest that VINE acts via Msb3 and the Msb3 adaptor BLOC-1 to inactivate Vps21. As Msb3 has not been reported to be regulated by phosphorylation, we reasoned that phosphorylation of its interactors could modulate Msb3 endosomal targeting or activity. The hexameric BLOC-1 complex, related to the human BLOC-1-related complex (BORC) (More et al., 2024) is required for Msb3 recruitment to endosomes but is dispensable for Msb3 GAP activity (John Peter et al., 2013; Rana et al., 2015). We used AlphaFold2 (Jumper et al., 2021) to predict the structure of the Vps21 GAP Msb3 bound to the six subunits of BLOC-1. In addition to an interaction between the C-terminal region of Msb3 and the BLOC-1 coiled-coil, this predicted a second, highly confident interaction involving residues 19-32 of the unstructured N-terminal domain of the BLOC-1 subunit Kxd1 which bound to Msb3 near, but not overlapping, its catalytic site (Figure 5A-B, Figure S5A).

**Figure 5.**
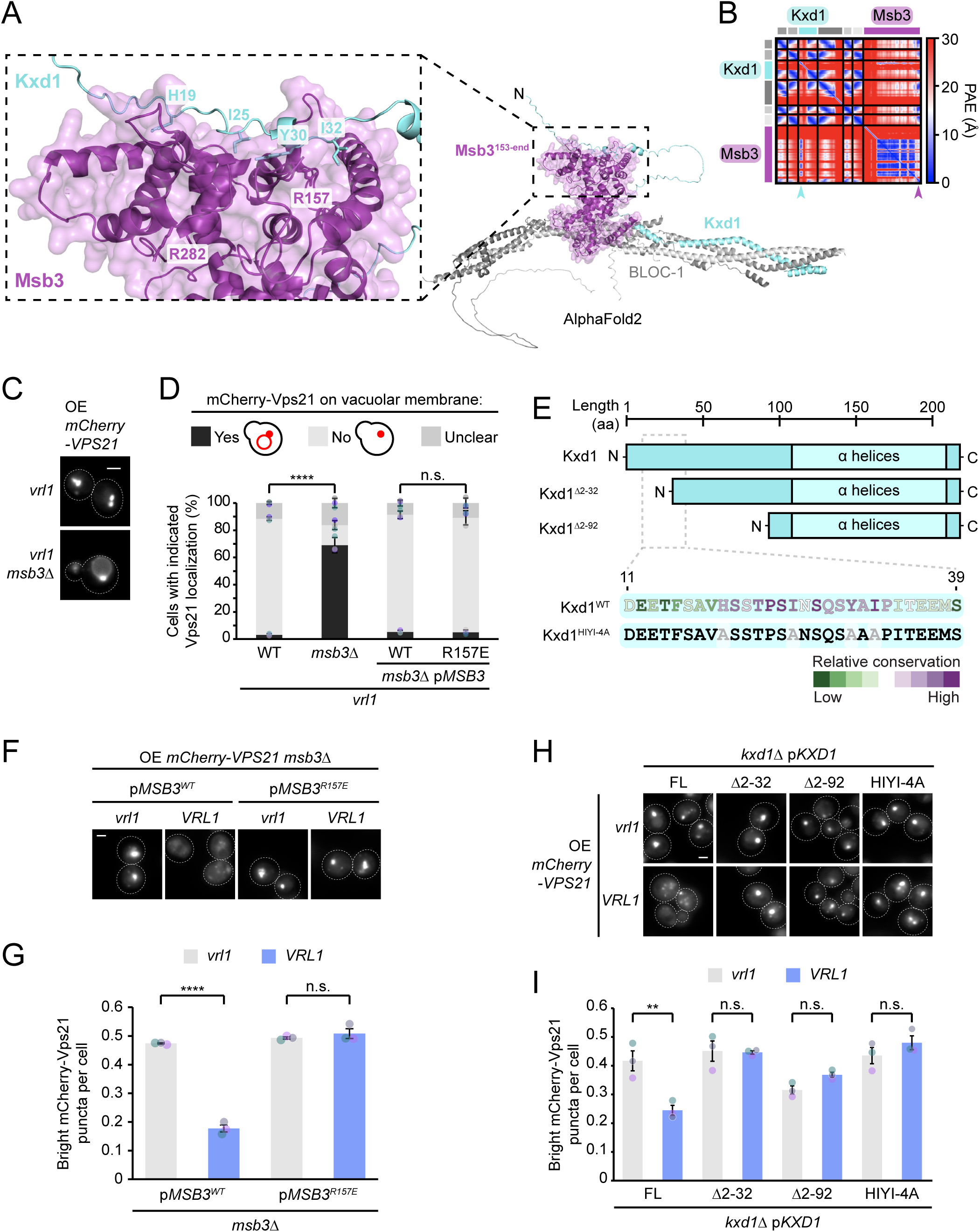
Regulatory sites in Msb3 and the BLOC-1 subunit Kxd1 mediate VINE-dependent Vps21 inactivation. (A) Msb3 and BLOC-1 are predicted to interact via two distinct interfaces. AlphaFold2-generated model showing predicted interaction of BLOC-1 subunits Bli1, Bls1, Kxd1, Vab2, Snn1 and Cnl1 with Msb3. The unstructured N-terminal region of Msb3 (residues 1-152) is not shown. Select residues in predicted interface and R282 of the Msb3 catalytic site are labelled in inset. (B) Plot showing PAE for predicted interaction in *A*. Arrows indicate region of Kxd1^Nt^ predicted to bind Msb3 (aqua) and C-terminal region in Msb3 predicted to bind to BLOC-1 coiled-coil (magenta). (C) Deletion of *MSB3* causes Vps21 to mislocalize to the vacuolar limiting membrane. Fluorescence micrographs of mCherry-Vps21-overexpressing yeast showing effect of *MSB3* deletion. (D) The predicted Kxd1^Nt^-interacting site in Msb3 is dispensable for Vps21 inactivation by VINE. Effect of *MSB3* deletion and complementation by plasmid-expressed *MSB3* alleles on RFP-Vps21 localization. Key shows strategy for manual quantitation of blinded images. One-way ANOVA with Tukey’s multiple comparisons test; n=3, cells/strain/replicate ≥ 78; n.s. = p > 0.05, **** = p < 0.0001. (E) Schematic of N-terminal truncations of Kxd1. Inset shows region predicted to interact with Msb3 with residues coloured by conservation among fungal homologs. (F) The predicted Kxd1^Nt^-interacting site in Msb3 is essential for Vps21 inactivation by VINE. Fluorescence micrographs of *msb3*Δ cells co-expressing mCherry-Vps21 from the *TEF2* promoter and untagged *MSB3* alleles from a plasmid, in the presence and absence of *VRL1*. (G) Automated quantitation of bright mCherry-Vps21 puncta from *F*. One-way ANOVA with Tukey’s multiple comparisons test; n=3, cells/strain/replicate ≥ 869; n.s. = p > 0.05, **** = p < 0.0001. (H) The predicted Msb3-interacting site in the Kxd1^Nt^ is essential for Vps21 inactivation by VINE. Fluorescence micrographs of *kxd1*Δ cells co-expressing mCherry-Vps21 from the *TEF2* promoter and Venus-tagged *KXD1* alleles from a plasmid, in the presence and absence of *VRL1*. (I) Automated quantitation of bright mCherry-Vps21 puncta in *H*. One-way ANOVA with Tukey’s multiple comparisons test; n=3, cells/strain/replicate ≥ 1052; n.s. = p > 0.05, ** = p < 0.01. Scale bars, 2 µm. Error bars report SEM. PAE, predicted aligned error. OE, overexpressed. HIYI-4A, H19A I25A Y30A I32A. FL, full-length. pTM, predicted TM. ipTM, interface predicted TM.

We used targeted mutagenesis to test the functional relevance of this secondary Kxd1^Nt^-Msb3 interaction. The complete loss of Kxd1 or Msb3 caused accumulation of Vps21 on vacuolar membranes as previously reported (Lachmann et al., 2012; John Peter et al., 2013), and this was independent of VINE (Figure 5C, Figure S5B). We found that a point mutation in Msb3 that was designed to specifically disrupt the secondary Kxd1^Nt^-Msb3 interface (Msb3^R157E^) had no effect on the VINE-independent function of Msb3 (Figure 5D). Similarly, Kxd1 alleles with N-terminal truncations or alanine substitutions in the predicted Msb3-binding site (Figure 5E) did not impair the VINE-independent function of BLOC-1 (Figure S5C). This indicates that the predicted interface between the Kxd1^Nt^ and Msb3 is not required for basal levels of Vps21 inactivation in cells that lack VINE.

In contrast, disrupting the interface between Msb3 and the Kxd1^Nt^ prevented Vps21 regulation by VINE. The Msb3^R157E^ mutation caused Vps21 to remain at bright puncta regardless of *VRL1* expression (Figure 5F, G). Similarly, truncating or mutating the Kxd1 N-terminal domain completely blocked the ability of VINE to inactivate Vps21 (Figure 5H, I). These results suggest that the unstructured N-terminal domain of Kxd1 creates a secondary interface between BLOC-1 and Msb3 that enhances Msb3 function, either through stabilizing GAP recruitment or increasing its activity. Because these sites were dispensable for the VINE-independent function of BLOC-1 and Msb3 but critical for Vps21 inactivation downstream of VINE this secondary interface appears to act exclusively in a VINE-dependent manner.

### Phosphorylation sites in the Kxd1 N-terminal domain determine Vps21 inactivation

Because our results suggest that an interaction between Msb3 and the Kxd1 N-terminal domain is required for VINE-dependent Vps21 regulation, we hypothesized that this secondary interaction is promoted by the VINE-Glc7 complex. The region of Kxd1 predicted to associate with Msb3 contains four conserved serine and threonine residues (Figure 6A), at least two of which are phosphorylated (Leutert et al., 2023). If phosphorylation of these sites regulates formation of the Kxd1^Nt^-Msb3 interface, blocking or mimicking phosphorylation of these residues should influence Vps21 distribution. Indeed, mutating these residues to alanine to prevent their phosphorylation caused overexpressed mCherry-Vps21 to localize to small, dispersed puncta (Figure 6B, C), phenocopying *VRL1* expression or *MSB3* overexpression (Figure 2I, J) and suggesting that the Kxd1^Nt^ promotes GAP function when dephosphorylated. Accordingly, this Kxd1 phosphorylation-deficient mutant rescued CPY secretion induced by overexpressed Vps21 (Figure 6D, E). Conversely, mutating the putative phosphorylation sites in Kxd1 to aspartate and glutamate to mimic constitutive phosphorylation prevented VINE from inactivating Vps21 (Figure 6B, C), suggesting phosphorylation of the Kxd1 N-terminus renders this site unable to regulate Msb3. These results suggest that dephosphorylation of the Kxd1 N-terminus promotes its binding to Msb3, which enhances GAP activity on Vps21.

**Figure 6.**
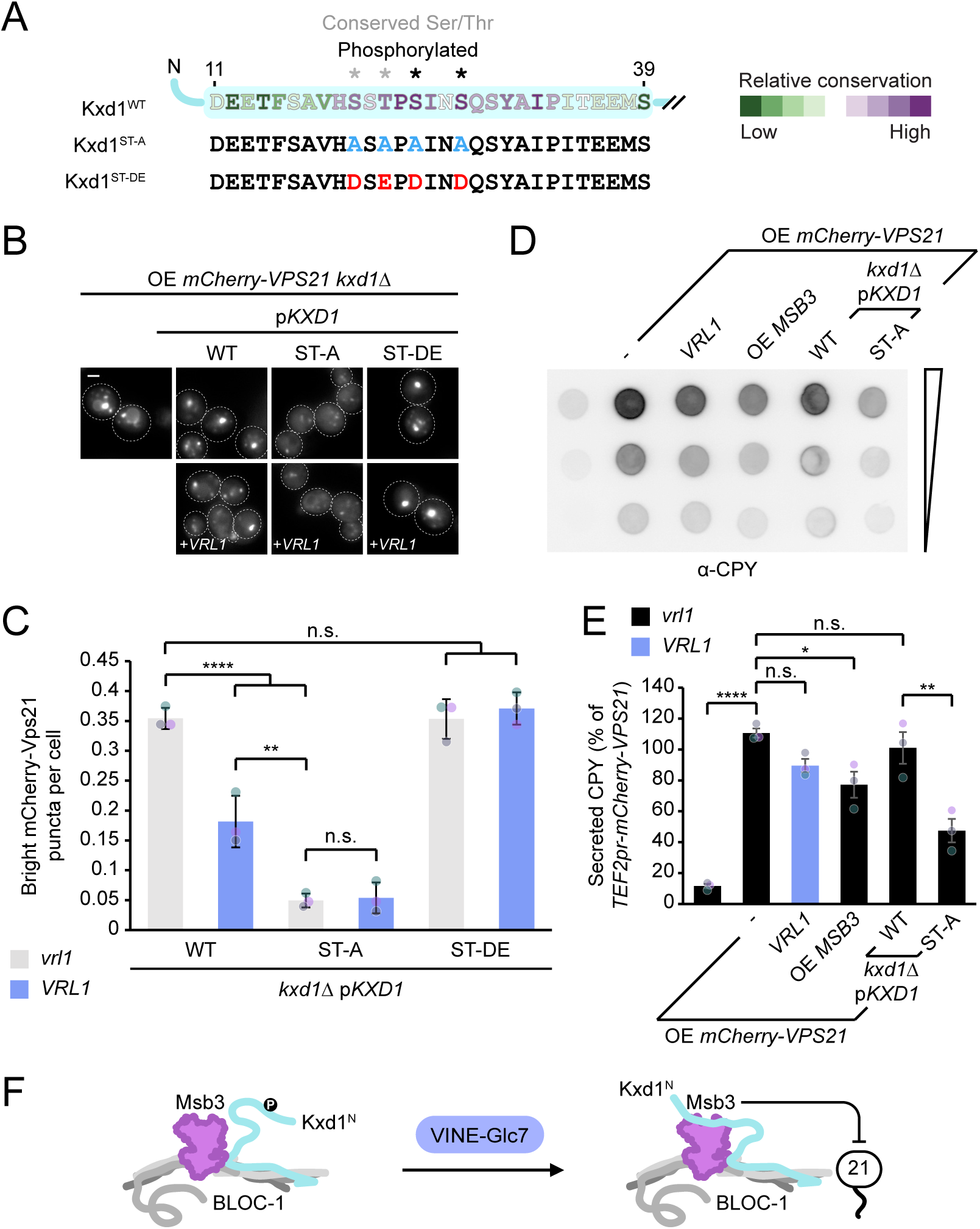
Phosphorylation sites in the Kxd1 N-terminal domain determine Vps21 inactivation. (A) Region of Kxd1 predicted to interact with Msb3, and schematic of phosphodeficient and phosphomimetic mutant alleles. Conserved serine and threonine residues in predicted interface are indicated with grey asterisks, and sites with previously detected phosphorylation (Leutert et al. 2023) are indicated with black asterisks. (B) Phosphorylation of the Kxd1^Nt^ determines Vps21 regulation. Fluorescence micrographs of *kxd1*Δ cells overexpressing mCherry-Vps21 and expressing *KXD1* WT or phospho-mutant forms from a plasmid, with or without *VRL1* expression. (C) Automated quantitation of bright mCherry-Vps21 puncta in *B*. One-way ANOVA with Tukey’s multiple comparisons test; n=3, cells/strain/replicate ≥ 1470; n.s. = p > 0.05, ** = p < 0.01. (D) Missorting of vacuolar cargo observed upon *VPS21* overexpression is rescued by Msb3 overexpression or by loss of Kxd1 phosphorylation. Western blot detecting CPY secreted from strains overexpressing mCherry-Vps21 showing effects of correction of the *vrl1* mutation by CRISPR, overexpression of GFP-Msb3 from the *NOP1* promoter, or expression of the Kxd1 phosphodeficient mutant. (E) Quantitation of secreted CPY in *D*. One-way ANOVA with Tukey’s multiple comparisons test; n=3; n.s. = p > 0.05, * = p < 0.05, ** = p < 0.01, **** = p < 0.0001. (F) Working model for VINE-dependent Vps21-inactivating pathway: phosphorylation of a regulatory site in the Kxd1 N-terminal domain prevents association with Msb3 and limits GAP-mediated inactivation of Vps21. A VINE-Glc7 complex promotes dephosphorylation of Kxd1, allowing its interaction with Msb3 and accelerating Vps21 inactivation. Scale bars, 2 µm. Error bars report SEM. Ser, serine. Thr, threonine. ST-A, S20A T22A S24A S27A. ST-DE, S20D T22E S24D S27D. OE, overexpressed. CPY, carboxypeptidase Y. GAP, GTPase-activating protein.

Together our data are consistent with a model in which the VINE-Glc7 complex directly or indirectly promotes dephosphorylation of the Kxd1 N-terminal domain to enhance the BLOC-1-dependent recruitment or activation of Msb3, accelerating Vps21 inactivation (Figure 6F).

## Discussion

We have identified a unique role for the VINE complex in regulating endosomal Rab GTPases. Whereas VINE activates Rab5-family proteins similarly to other VPS9-family GEFs, we report that it also orchestrates the inactivation of the yeast Rab5 homolog Vps21. Our data support a model in which the ankyrin repeat-containing domain of Vrl1 recruits the phosphatase Glc7, enabling an interaction between the GAP Msb3 and a regulatory site in the BLOC-1 subunit Kxd1. This interaction stimulates GAP-mediated inactivation of Vps21. Because VINE both promotes Vps21 inactivation and catalyzes nucleotide exchange on other Rab5-family GTPases, it has the capacity to suppress the function of one endosomal Rab while enhancing the activity of others.

### VINE is a novel endosomal PP1 adaptor

Our findings indicate that VINE functions as a novel endosomal adaptor for the PP1 catalytic subunit Glc7. Glc7 associates with a conserved surface on the Vrl1 ankyrin repeat-containing domain. Although Glc7 has more than 20 known binding partners including both activators and inhibitors, the regulatory consequences of many of these interactions remain unclear (Cannon, 2010). Our experiments suggest that, when recruited to VINE, Glc7 promotes dephosphorylation of the unstructured N-terminal region of Kxd1. The most straightforward interpretation is that VINE recruits catalytically active Glc7, which directly dephosphorylates Kxd1. Consistent with this model, structural predictions indicate that VINE binding does not occlude the Glc7 active site, suggesting that VINE acts as a positive regulator of Glc7. However, we cannot exclude the possibility that Glc7 acts indirectly, for example by inactivating kinases that target Kxd1.

Glc7 has well-established VINE-independent roles in endocytic trafficking and vacuole fusion (Peters et al., 1999; Bryant and James, 2003). This suggests it may already be present on endosomes, and that VINE acts to couple Glc7 activity to specific endosomal substrates. It will be important in future studies to determine whether recruitment of Glc7 by VINE also influences additional downstream events at endosomes or vacuoles.

It also remains unclear whether VINE primarily positions Glc7 near substrates or enhances Glc7 catalytic activity. Notably, the VINE-Glc7 interactions may be spatially and temporally restricted. We detected Glc7 association with Vrl1 at endosomes only using the sensitive split-Venus assay, suggesting that not all VINE complexes necessarily contain Glc7. Phosphoregulation of Kxd1 may therefore take place in specific endosomal subdomains or at a specific stage of endosome-to-vacuole transport. Such spatial restriction could explain why VINE selectively affects only a subset of Vps21 effectors.

### A novel regulatory mechanism for enhancing GAP activity

Our work identifies a regulatory site within the N-terminal region of the BLOC-1 subunit Kxd1 as a key target of the VINE-Glc7 pathway. While we cannot exclude the possibility that VINE-Glc7 acts on additional substrates, our data suggest that dephosphorylation of the Kxd1 N-terminal domain is both required and sufficient to account for VINE-dependent regulation of Vps21. This supports the model that phosphorylation of the Kxd1 N-terminus inhibits BLOC-1- and Msb3-dependent Vps21 inactivation, and that VINE-Glc7 relieves this inhibition.

We propose that Msb3 engages the BLOC-1 complex through two functionally distinct interactions: whereas binding to the central coiled-coil region of BLOC-1 mediates basal recruitment of Msb3 to endosomes (John Peter et al., 2013), interaction with the unstructured Kxd1 N-terminal domain enhances Msb3 function. The mechanism by which interaction with the Kxd1 N-terminus enhances Msb3 activity remains unclear. One possibility is that binding to the dephosphorylated Kxd1 tail increases the stability or residence time of Msb3 at endosomes.

In addition to endosomes, Msb3 localizes to other cellular sites including regions of polarized growth where it is recruited by the polarisome component Spa2 (Tcheperegrine et al., 2005; Bareis et al., 2026). If these adaptors compete dynamically for Msb3 binding, a bipartite interaction with BLOC-1 could bias Msb3 localization toward endosomes. Consistent with this idea, endosomal Msb3 is difficult to detect under conditions where BLOC-1 subunits are readily visible, suggesting that BLOC-1/Msb3 interactions are substoichiometric (John Peter et al., 2013). Increased contacts between BLOC-1 and Msb3 could therefore stabilize Msb3 recruitment to endosomes and accelerate Vps21 inactivation.

Alternatively, the interaction between dephosphorylated Kxd1 and Msb3 could induce a conformational change that enhances GAP catalytic activity. Future *in vitro* assays will be required to test this possibility. Although regulation of GAP activity through such mechanisms is unusual, there is precedent for phosphoregulation of Rab regulators. For example, phosphorylation alters the activity and Rab specificity of the Rab43-specific GAP USP6NL (Lanzetti et al., 2007; Wang et al., 2026). Phosphorylation of unstructured regions also controls the recruitment and function of several Rab regulators (Mafakheri et al., 2018; Eickelschulte et al., 2021; Kim et al., 2019; Locke and Thorner, 2019). Together, these examples support the idea that phosphoregulation of flexible protein regions can dynamically tune Rab regulatory activity.

### Specialized roles of VPS9-family GEFs

Our findings further demonstrate that VINE has functions not shared by other VPS9-family GEFs. Although Muk1, Vps9, and Vrl1 have partially redundant roles in protein trafficking (Paulsel et al., 2013; Cabrera et al., 2012; Locke and Thorner, 2019; Shortill et al., 2022; Li et al., 2019), we found that VINE and Vps9 can oppose one another in specific contexts, consistent with previous data indicating that individual VPS9-family members differ in their contributions to endosomal processes (Shideler et al., 2015).

VINE and Vps9 represent distinct, ancient VPS9-family GEF lineages containing VARP and Rabex-5 respectively (Herman et al., 2018). The loss of Vrl1/VARP homologs in multiple lineages suggests that this GEF family may play specialized roles that are important only under particular physiological or stress conditions (Herman et al., 2018). The absence of obvious fitness defects upon loss of VINE in laboratory strains is consistent with this. Despite the absence of VARP- and VINE-like GEFs in some organisms, the broad conservation of these families suggests that they fulfill complementary functions in endosomal regulation. In fact, the Vps9-like protein Rabex-5 has been proposed to maintain Rab activation until ubiquitylated cargo is sequestered into intralumenal vesicles (Mattera and Bonifacino, 2008; Ott et al., 2025). The presence of a ubiquitin-binding CUE domain in yeast Vps9 suggests that this regulatory strategy may be conserved (Davies et al., 2003; Shideler et al., 2015).

Endosomes are tubulo-vesicular organelles, with vesicular domains where intralumenal vesicles form separated from tubular domains that mediate cargo recycling. Vps9 may function primarily at vesicular regions involved in anterograde cargo sorting, whereas VINE, through its PX-BAR dimer, targets PI3P-rich endosomal membranes and may generate retrograde sorting tubules similar to those formed by retromer (Shortill et al., 2022). We propose that the distinct and sometimes opposing functions of these GEFs reflect their activity at spatially distinct endosomal subdomains.

### Dual Rab regulation by VINE and retromer

The VINE-dependent pathway of Vps21 inactivation raises the question of why a Rab GEF would both promote and oppose endosomal Rab signaling. One possibility is that VINE buffers the local equilibrium between Vps21-GTP and Vps21-GDP, limiting excessive Rab activity adjacent to regions containing other VPS9-family GEFs or under conditions of elevated Rab expression. Notably, our findings predict that Vps21 inactivation by VINE is restricted to membrane domains containing BLOC-1.

VINE exhibits notable parallels with the retromer complex, which targets membranes through sorting nexin (SNX) adaptors and recruits both Rab GEFs and GAPs, forming a hub for endosomal Rab regulation (Hesketh et al., 2014; Seaman et al., 2009; Jia et al., 2016; Antón-Plágaro et al., 2025). Through these interactions, retromer has been proposed to facilitate Rab switching, coupling inactivation of an upstream Rab to activation of a downstream Rab at defined membrane subdomains (Antón-Plágaro et al., 2025). Similarly, VINE is an endosomal SNX-containing complex (Shortill et al., 2022) that acts as a GEF while promoting GAP function. VINE therefore has the potential to bias Rab signaling toward minor Rab5 isoforms such as Ypt52 in a context-dependent manner.

Together, these observations suggest that VINE represents a second endosomal Rab regulatory hub analogous to retromer. Whether the dual GEF and GAP-promoting activities of VINE are themselves regulated, or confined to specific endosomal subdomains, remains an important question for future study. More broadly, our findings demonstrate that like retromer, VINE has the potential to spatially and temporally organize Rab signaling during endosomal trafficking and maturation.

## Materials and Methods

| Key Resources Table |  |  |  |  |
| --- | --- | --- | --- | --- |
| Reagent type (species) or resource | Designation | Source or reference | Identifiers | Additional information |
| Yeast strain collection | Schuldiner SWAT NOP1pr-GFP-Collection | Weill et al., 2018 |  |  |
| Antibody | Anti-CPY (Mouse monoclonal) | Thermo Fisher Scientific | A-6428 | 1:500 |
| Antibody | Anti-HA (Mouse monoclonal) | Sigma-Aldrich | H9658; HA-7 | WB (1:1000) |
| Antibody | Anti-GFP (Mouse monoclonal) | Roche | 11-814-460-001 | WB (1:1000) |
| Antibody | Anti-GFP (Rabbit polyclonal) | Eusera | EU2 | CoIP |
| Antibody | HRP-Anti-Mouse (Goat polyclonal) | Jackson | 115-035-146 | WB (1:20000) |
| Other | nProtein A Sepharose 4 Fast Flow | Cytiva | 17528004 | CoIP |
| Other | Amersham ECL | Cytiva | GERPN2209 | Chemiluminescent reagent |
| Other | Yeast/Fungal ProteaseArrest | G-Biosciences | 786-435 | Yeast protease inhibitor; 100X |
| Other | Concanavalin A | Sigma-Aldrich | C2010 | Yeast live-cell imaging preparation |
| Other | FM4-64 | Invitrogen | T3166 | Vacuolar Rim Stain; 4 μM |
| Other | CMAC | Setareh Biotech | 6627 | Vacuole Stain; 100 μM |
| Chemical compound, drug | Methotrexate | Sigma Aldrich | PHR1396 | DHFR inhibitor; 200 μg/ml, DMSO |
| Software, algorithm | CellProfiler | Lamprecht et al., 2007 |  | Yeast colony array image analysis |
| Software, algorithm | MetaMorph | MDS Analytical Technologies | Version 7.8.13.0 | Automated image analysis |
| Software, algorithm | GraphPad Prism | GraphPad Software | Version 10.2.2 | Statistical analysis |
| Software, algorithm | ImageJ | NIH |  | Western blot densitometry |
| Software, algorithm | ColabFold AlphaFold2 Advanced | Jumper et al., 2021; Mirdita et al., 2022 |  | Protein structure and binding prediction software |
| Software, algorithm | AlphaFold3 Server | Abramson et al. 2024 |  | Protein structure and binding prediction software |
| Software, algorithm | ConSurf | Ashkenazy et al., 2016 |  | Protein conservation analysis |

### Yeast strains and plasmids

Yeast strains and plasmids used in this study are detailed in Tables S2 and S3. Genomic integration of recombinant DNA was carried out by homologous recombination or by Cas9-mediated integration (Bean et al. 2022). Genetic manipulations were confirmed by colony PCR, microscopy and western blotting as appropriate. Plasmids were constructed in specified pRS vectors using NEBuilder (New England BioLabs, Ipswich, Massachusetts) in *Escherichia coli* or by homologous recombination in yeast and confirmed by DNA sequencing or constructed using Golden Gate cloning in *E. coli*. *VRL1* was expressed either from its endogenous promoter, or from the stronger *ADH1* promoter. Both promoters result in the formation of similar levels of VINE, as Vin1 levels become limiting when only Vrl1 is overexpressed (Shortill et al. 2022).

### Fluorescence microscopy and automated image analysis

Yeast strains for microscopy were cultured overnight in minimal media excluding amino acids for selective growth of plasmid-containing strains as appropriate. Culture media was replaced four hours before log-phase cells were plated on 96-well glass-bottom plates (Cellvis, Mountain View, California) treated with concanavalin A. Vacuoles were stained with FM4-64 (Invitrogen, Waltham, Massachusetts) or CMAC (Setareh Biotech, San Jose, California) for 30 minutes, after which cells were rinsed and incubated at 30°C for one hour before imaging. Cells were imaged with Leica DMi8 microscope (Leica Microsystems, Wetzlar, Germany) with an ORCA-flash 4.0 digital camera (Hamamatsu Photonics, Shizuoka, Japan) and a Plan-Apochromat 63x/1.30 glycerol immersion lens (Leica Microsystems, Wetzlar, Germany). MetaMorph 7.8.13.0 (MDC Analytical Technologies, Sunnyvale, California) was used to acquire and process images. Linear intensity scale changes were uniformly applied to all images of a given fluorophore within an experiment. Where appropriate this common scaling is shown in an inset and the full-size image is scaled to clearly show relevant morphological features. Images were exported via Photoshop CC 2025 (Adobe, San Jose, California) to Illustrator CC 2021 (Adobe, San Jose, California). Automated quantitation was performed on unmodified images using MetaMorph 7.8.13.0 journals (MDC Analytical Technologies, Sunnyvale, California). Dead cells were masked and live cells identified using the Count Nuclei feature, and puncta identified using size and intensity above local background (IALB) as criteria in the Granularity feature. To generate heatmaps summarizing image quantitation data, Microsoft Excel 2025 (Microsoft, Redmond, Washington) was used to place average values on a colour gradient from blue (0 large bright puncta per cell) to black (the number of puncta per cell in the corresponding control strain with endogenous levels of Vps9 and Muk1 and no Vrl1) to yellow (the maximum number of puncta per cell observed for the given GTPase). Manual quantitation of uniformly scaled blinded images was performed in ImageJ (Schneider et al. 2012) using the Multi-Point tool.

### Detection of secreted CPY-Invertase

Secreted invertase was detected as described in Darsow et al. (2000) and Dalton et al. (2015). Yeast strains were cultured overnight in synthetic media containing 2% fructose and lacking uracil. 10 OD_600_ equivalents of culture were collected, washed in media and 20 μL added to 30 μL of 0.1 M sodium acetate pH 4.9. 12.5 μL of 0.5 M sucrose was added to samples and glucose standards and incubated at 30°C for 30 minutes. Invertase was inactivated by adding 75 μL of 0.2 M pH 10 potassium phosphate and boiling for 5 minutes. Samples and standards were then incubated for 30 minutes at 30°C with 500 μL of glucostat solution (0.1 M pH 7 potassium phosphate, 2 U/mL glucose oxidase, 2.5 µg/mL horse radish peroxidase, 12.5 µg/mL N-ethyl maleimide and 150 µg/mL o-dianisidine). Reactions were arrested by adding 500 μL of 6N HCl. Absorbance at 540nm was measured in a 96-well microtiter plate (167008, Thermo Fisher Scientific) using the Spark Multimode Microplate Reader (Tecan, Männedorf, Switzerland). Units of invertase activity indicate the amount of glucose (in nmol) produced by one OD_600_ of yeast each minute.

### Detection of extracellular CPY by western blot

Yeast strains were cultured overnight in minimal amino acid dropout media selective for plasmid-bearing cells. Strains were diluted to a uniform optical density at 600nm (OD_600_) and grown for four hours or to an OD_600_ of over 1.0. Cultures were diluted to OD_600_ of 1 in media, serially diluted 1:1 with sterile water, and spotted onto synthetic media. Plates were overlaid with a nitrocellulose membrane and incubated for 12 hours at 30°C. Positive control strains secreting a high level of CPY were spotted in duplicate to provide separate data points for normalization and statistical comparison. Membranes were rinsed and probed with mouse anti-CPY antibody (A-6428, Thermo Fisher Scientific) and HRP-conjugated goat anti-mouse polyclonal antibody (115–035-146; Jackson ImmunoResearch Laboratories) and chemiluminescence signal measured using the Vilber Fusion FX (Vilber Smart Imaging, Collégien, France).

### Prediction of protein folding and complex formation and bioinformatic analysis of sequence conservation

Predictive structural models of proteins, protein fragments or multiprotein complexes were generated using the ColabFold AlphaFold2 advanced server (Jumper et al. 2021, Mirdita et al. 2022) with default settings or using the AlphaFold3 server (Abramson et al. 2024). Amino acid conservation was projected onto protein sequences or predictive structural models using ConSurf (https://consurf.tau.ac.il; Ashkenazy et al., 2016) with default settings. Predicted structural models were prepared for presentation using PyMOL (Schrödinger, LLC, New York, New York).

### Yeast spheroplasting and coimmunoprecipitation

Yeast strains were cultured in minimal selective media to log phase and 75 OD_600_ equivalents of cells harvested. After incubating for 15 minutes at room temperature in 50 mM Tris-Cl with 10 mM DTT pH 9.5, cells were rinsed and cell walls digested using spheroplasting buffer (2 M Sorbitol 1 M KH_2_PO_4_ 1 M MgCl_2_ 250 μg/ml zymolase, pH 7.4) for 1 hour at 30°C. Spheroplasts were washed twice using 1.2 M sorbitol and frozen at −80°C. Thawed spheroplasts were lysed in 50 mM NaCl 1 mM EDTA 50 mM HEPES 0.2% Tween 20 1 mM PMSF (pH 7.4) with protease inhibitors (G Biosciences). 50 μl of cleared lysate was reserved and combined with 50 μl of 2X Laemmli sample buffer (4% SDS, 20% glycerol, 120 mM Tris-Cl (pH 6.8), 0.01 g bromophenol blue and 10% beta-mercaptoethanol). Remaining cleared lysate was incubated for one hour with 4 μl rabbit anti-GFP antibody (EU2, Eusera) then for one hour with protein A sepharose beads (Cytiva). Beads were washed three times in 50 mM NaCl 1 mM EDTA 50 mM HEPES 0.2% Tween 20 (pH 7.4) and samples eluted in Thorner buffer (8 M Urea, 5% SDS, 50 mM Tris-Cl (pH 6.8), 1% beta-mercaptoethanol and 0.4 mg/mL bromophenol blue) by heating to 80°C for 5 minutes. Samples were separated on 10% SDS-PAGE gels and epitope-tagged proteins detected using mouse anti-HA (H9658, Clone HA-7, Sigma-Aldrich) or mouse anti-GFP (11-814-460-001, Roche) antibodies followed by goat anti-mouse polyclonal antibody conjugated to horseradish peroxidase (HRP) (115-035-146, Jackson ImmunoResearch Laboratories). Active HRP was detected using Amersham ECL (GERPN2209, Cytiva) and Vilber Fusion FX (Vilber Smart Imaging, Collégien, France). Intensity of bands or secreted protein was measured using ImageJ (Schneider et al. 2012).

### DHFR protein fragment complementation assay

Bait strains were generated in the BY4742 background by transforming a *DHFR[1,2]-VRL1* plasmid and a *LaG16-DHFR[3]* prey adaptor plasmid expressing a fusion of the GFP-binding nanobody LaG16 (Fridy et al. 2014) to the C-terminal DHFR fragment. Using a BM3-BC robot (S&P Robotics Inc., Toronto, Canada), bait strains were arrayed and mated to the MATa SWAT collection expressing N-terminally GFP-tagged strains from the *NOP1* promoter (Yofe et al. 2016). Diploid strains harboring plasmids were selected by pinning to media containing Hygromycin B and lacking leucine and uracil. Diploid arrays were pinned to plates containing methotrexate and lacking methionine, lysine and leucine, cultured for one day at 30°C and pinned to a duplicate set of methotrexate-containing plates. Plates were imaged daily for 7 days and colony growth measured using CellProfiler (Lamprecht et al., 2007). Interaction Z-scores with wild type *VRL1-DHFR[1,2]* were generated using Microsoft Excel 2025 (Microsoft, Redmond, Washington) and for all preys with Z-score > 1.0, the colony area reporting the interaction with Vrl1^WT^ was divided by the colony area reporting the interaction with Vrl1^DN-NS^ and this ratio used to generate ratio Z-scores.

### Statistical analysis and graphical representation of quantitative data

Statistical tests were performed using GraphPad Prism 10.2.2 (GraphPad Software, San Diego, California) as described in figure legends. Normality of data was assumed but not formally tested. Hypotheses were tested against a threshold of 95% confidence (P < 0.05). Quantitative data were graphed using Microsoft Excel 2025 (Microsoft, Redmond, Washington). The mean of measurements from a minimum of 3 independent replicates is graphed as column charts, with data points from all individual replicates shown as overlaid scatter plots. Error bars indicate standard error of the mean.

## Supplemental Material

Figures S1-S5 contain the results of microscopy and co-immunoprecipitation experiments and predictive structural modeling. Table S1 contains the full dataset from the nanobody-adapted split-DHFR screen. Tables S2-S4 contain information describing yeast strains, plasmids and oligonucleotides used in this study, respectively.

## Data Availability Statement

All data generated in this study are included in the manuscript and supplemental files.

## Acknowledgements

We thank Dr. Maya Schuldiner (Weizmann Institute of Science, Rehovot, Israel) for sharing the SWAT yeast library, Drs. Bjorn Bean and Vincent Martin (Concordia University, Montreal, Canada) for sharing yeast CRISPR reagents, Dr. Alexey Merz (University of Washington, Seattle, USA) for sharing yeast strains, Dr. Greg Odorizzi (University of Colorado, Boulder, USA) and Dr. Christopher Loewen (University of British Columbia, Vancouver, Canada) for sharing plasmids, and Dr. Luc Berthiaume (University of Alberta, Edmonton, Canada) for sharing rabbit anti-GFP serum.

This work was supported by the Natural Sciences and Engineering Research Council of Canada (grants RGPIN-2022-04573 and RGPIN-2016-04290 to EC and PGS-D3 Doctoral Scholarship to SPS); Canada Foundation for Innovation (Leading Edge Fund 30636); Canadian Institutes for Health Research (grant PJT-180544 to EC, CGS-M Frederick Banting and Charles Best Canada Graduate Scholarship to SPS and MSF, and CGS-D Frederick Banting and Charles Best Canada Graduate Scholarship to MSF); BC Children’s Hospital Research Institute (Jan M Friedman Graduate Studentship to SPS); and the University of British Columbia (Affiliated Fellowships and 4-Year Doctoral Fellowship to SPS and MSF).

## Supplementary Figure and File Legends

**Figure S1.**
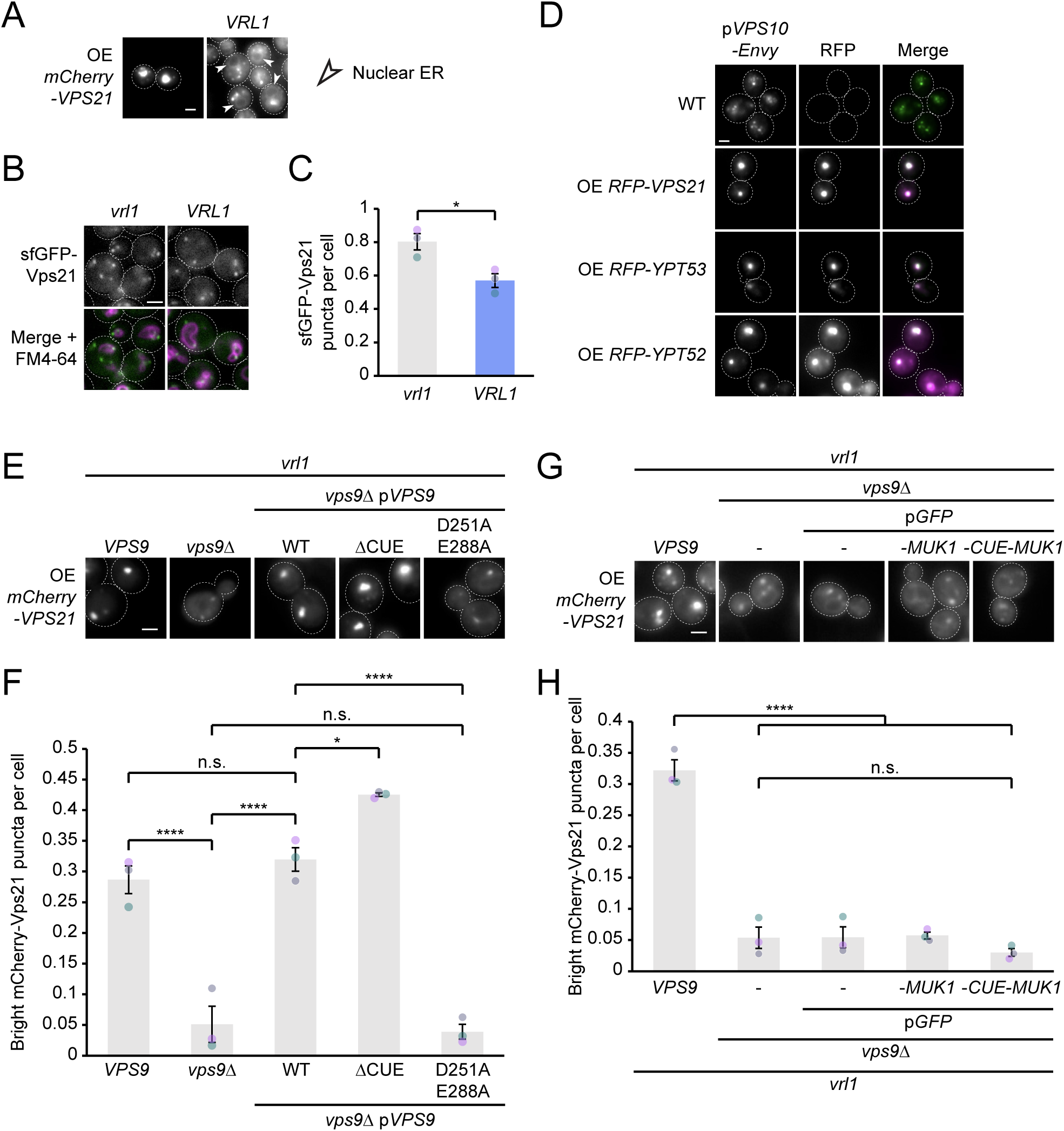
(A) VINE partially redistributes Vps21 to the nuclear ER. Fluorescence micrographs of cells overexpressing mCherry-Vps21 with or without untagged Vrl1 expressed from its endogenous promoter. Images scaled to show faint mCherry signal at ER, marked by white arrows. (B) VINE affects the localization of endogenously expressed Vps21. Fluorescence micrographs of cells expressing sfGFP-Vps21 from the *VPS21* promoter, with vacuoles stained by FM4-64. (C) Automated quantitation of bright sfGFP puncta per cell, in *B*. Two-tailed equal variance t test; n=3, cells/strain/replicate ≥ 2710; * = p < 0.05. (D) Vps10 colocalizes with overexpressed endosomal Rab proteins. Fluorescence micrographs of cells overexpressing mCherry-Rab5 fusions and expressing GFP-tagged CPY receptor Vps10 from a plasmid. (E) Vps9 GEF activity drives endosomal morphology defects through overexpressed Vps21. Fluorescence micrographs of WT and *vps9Δ* cells co-overexpressing mCherry-Vps21 and plasmid-borne Vps9 alleles. (F) Quantitation of bright mCherry-Vps21 puncta in *E*. One-way ANOVA with Tukey’s multiple comparisons test; n=3, cells/strain/replicate ≥ 1385; n.s. = p > 0.05, * = p < 0.05, **** = p < 0.0001. (G) Fluorescence micrographs of WT and *vps9Δ* cells overexpressing mCherry-Vps21 and expressing GFP, GFP-Muk1 or a GFP-tagged fusion of the Vps9 CUE domain to Muk1 from plasmids. (H) Quantitation of bright mCherry-Vps21 puncta in *G*. One-way ANOVA with Tukey’s multiple comparisons test; n=3, cells/strain/replicate ≥ 1521; n.s. = p > 0.05, **** = p < 0.0001. Scale bars, 2 µm. Error bars report SEM. OE, overexpressed.

**Figure S2.**
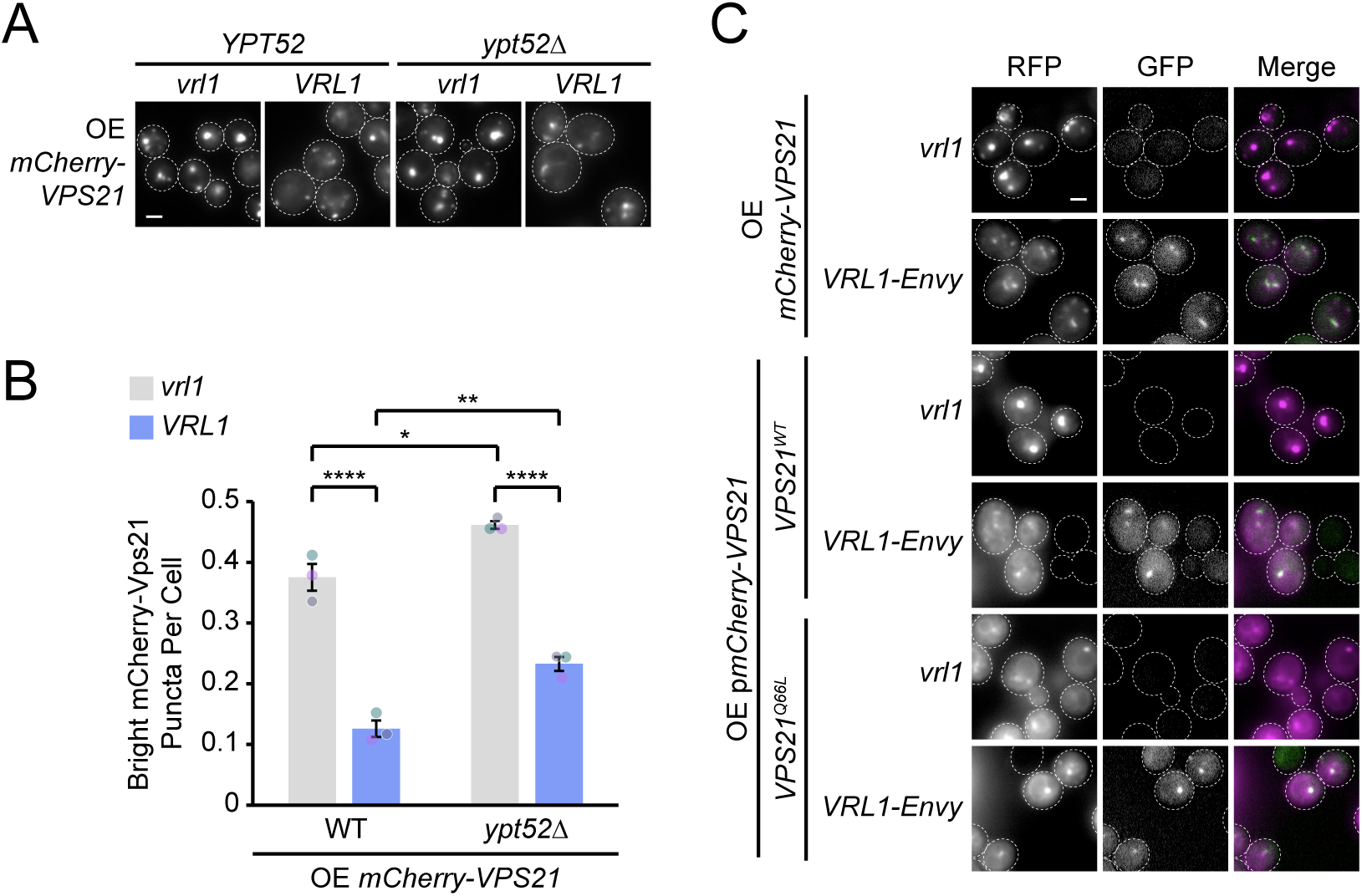
(A) VINE does not require Ypt52 to negatively regulate Vps21. Fluorescence micrographs of WT and *ypt52*Δ cells overexpressing mCherry-Vps21 with or without plasmid-expressed *VRL1*. (B) Quantitation of bright mCherry-Vps21 puncta in *A*. One-way ANOVA with Tukey’s multiple comparisons test; n=3, cells/strain/replicate ≥ 1293; * = p < 0.05, ** = p < 0.01, **** = p < 0.0001. (C) A mutation in Vps21 blocking GTP hydrolysis prevents its regulation by VINE. Fluorescence micrographs of cells overexpressing mCherry-Vps21, or the constitutively active Vps21^Q66L^ allele, with or without expression of GFP-tagged Vrl1. Scale bars, 2 µm. Error bars report SEM. OE, overexpressed.

**Figure S3.**
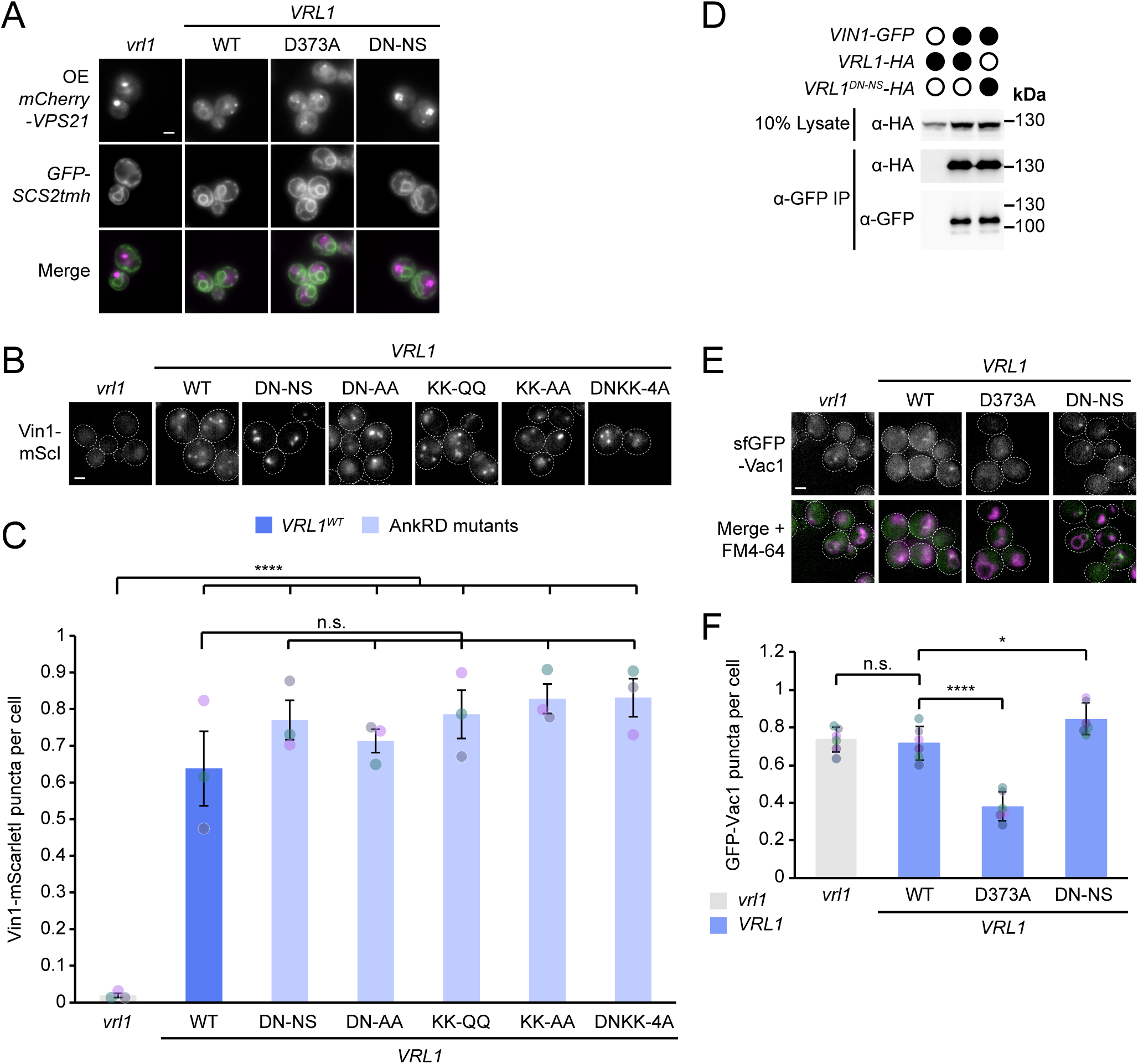
(A) The conserved site on the Vrl1 AnkRD is required to redistribute Vps21 to the nuclear ER. Fluorescence micrographs of cells overexpressing mCherry-Vps21 and expressing the ER marker Scs2tmh with or without untagged Vrl1 alleles expressed from the *VRL1* promoter. Images are scaled to show faint mCherry signal at ER. (B) The conserved site on the Vrl1 AnkRD is not required to localize Vin1 to endosomes. Fluorescence micrographs of cells expressing Vin1-mScI with or without plasmid-expressed WT Vrl1 or Vrl1 alleles with mutations in the AnkRD. (C) Quantitation of Vin1-mScI puncta in *B*. One-way ANOVA with Tukey’s multiple comparisons test; n=3, cells/strain/replicate ≥ 650; n.s. = p > 0.05, **** = p < 0.0001. (D) The novel conserved site on the Vrl1 AnkRD is not required to bind Vin1. Western blot showing immunoprecipitated Vin1-GFP and co-immunoprecipitated HA-tagged Vrl1 alleles. (E) The Vrl1 GEF active site and the conserved AnkRD site have opposite effects on Vps21 effector localization. Fluorescence micrographs of cells expressing sfGFP-Vac1 and untagged Vrl1 alleles, with vacuoles labelled by FM4-64. (F) Automated quantitation of GFP-Vac1 puncta in *E*. One-way ANOVA with Dunnett’s multiple comparisons test; n=6, cells/strain/replicate ≥ 1116; n.s. = p > 0.05, * = p < 0.05, **** = p < 0.0001. Scale bars, 2 µm. Error bars report SEM. *SCS2tmh*, transmembrane helix of *SCS2*. DN-NS, D638N N642S. DN-AA, D638A N642A. KK-QQ, K669Q K679Q. KK-AA, K669A K679A. DNKK-4A, D638A N642A K669A K679A. mScI, mScarletI. AnkRD, ankyrin repeat-containing domain. OE, overexpressed. IP, immunoprecipitate. sfGFP, superfolder GFP.

**Figure S4.**
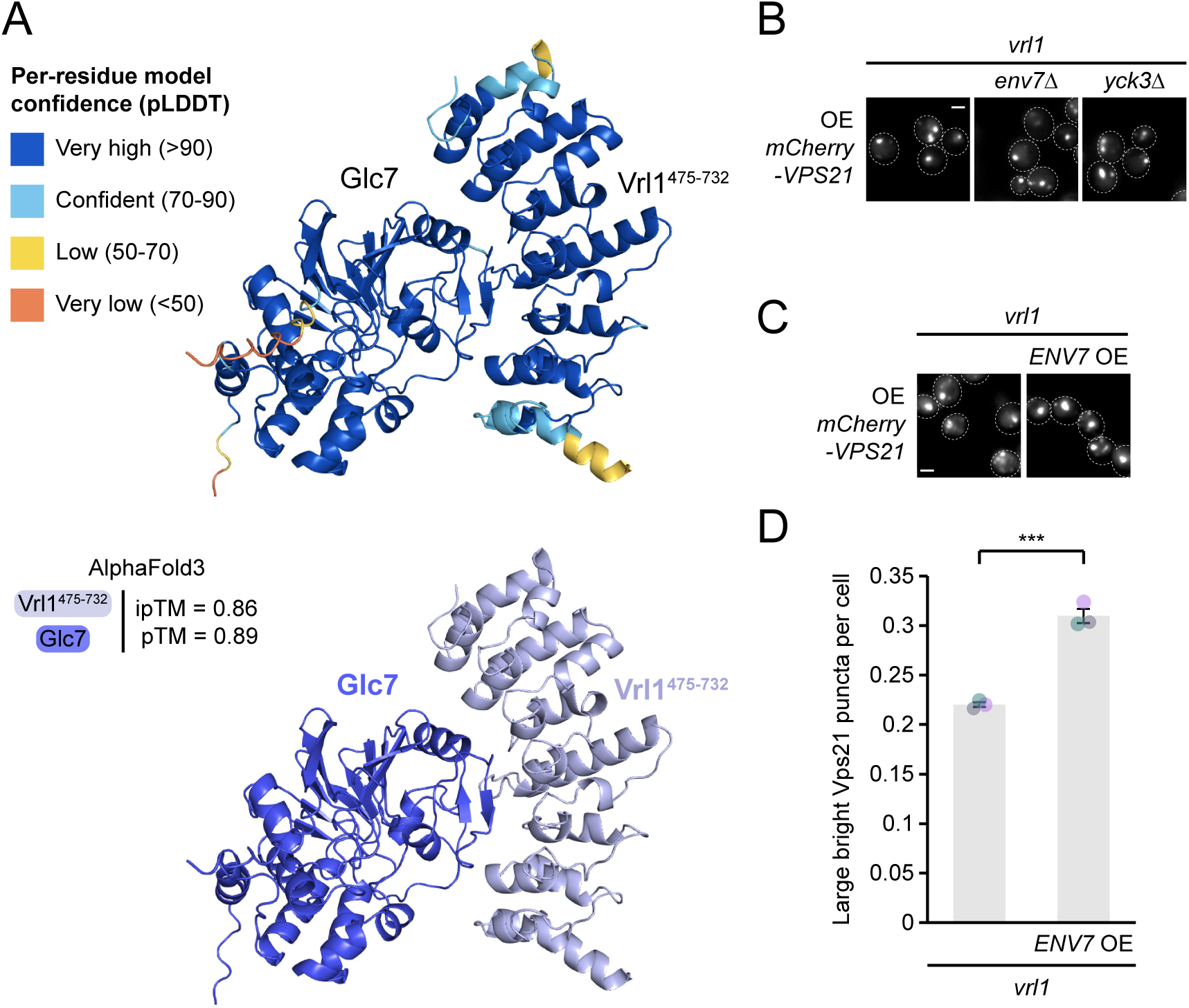
(A) AlphaFold3 confidently predicts an interaction between Glc7 and the Vrl1 AnkRD. AlphaFold3-predicted model of Vrl1 AnkRD with Glc7 coloured by pLDDT (top) and by chain (bottom). pTM and ipTM scores are indicated; sequences were trimmed to remove unstructured regions. (B) Vps21 overexpression drives endosomal defects independently of the kinases Env7 and Yck3. Fluorescence micrographs of the indicated WT and mutant cells expressing mCherry-Vps21 from the strong *TEF2* promoter. (C) Env7 overexpression exacerbates endosomal defects driven by Vps21 overexpression. Fluorescence micrographs of cells overexpressing mCherry-Vps21 showing effect of overexpressing untagged Env7. (D) Automated quantitation of large bright mCherry-Vps21 puncta in *C*. Two-tailed equal variance t test; n=3, cells/strain/replicate ≥ 1388; *** = p < 0.001. Scale bars, 2 µm. Error bars report SEM. OE, overexpressed. pLDDT, predicted local-difference distance test. pTM, predicted TM. ipTM, interface predicted TM.

**Figure S5.**
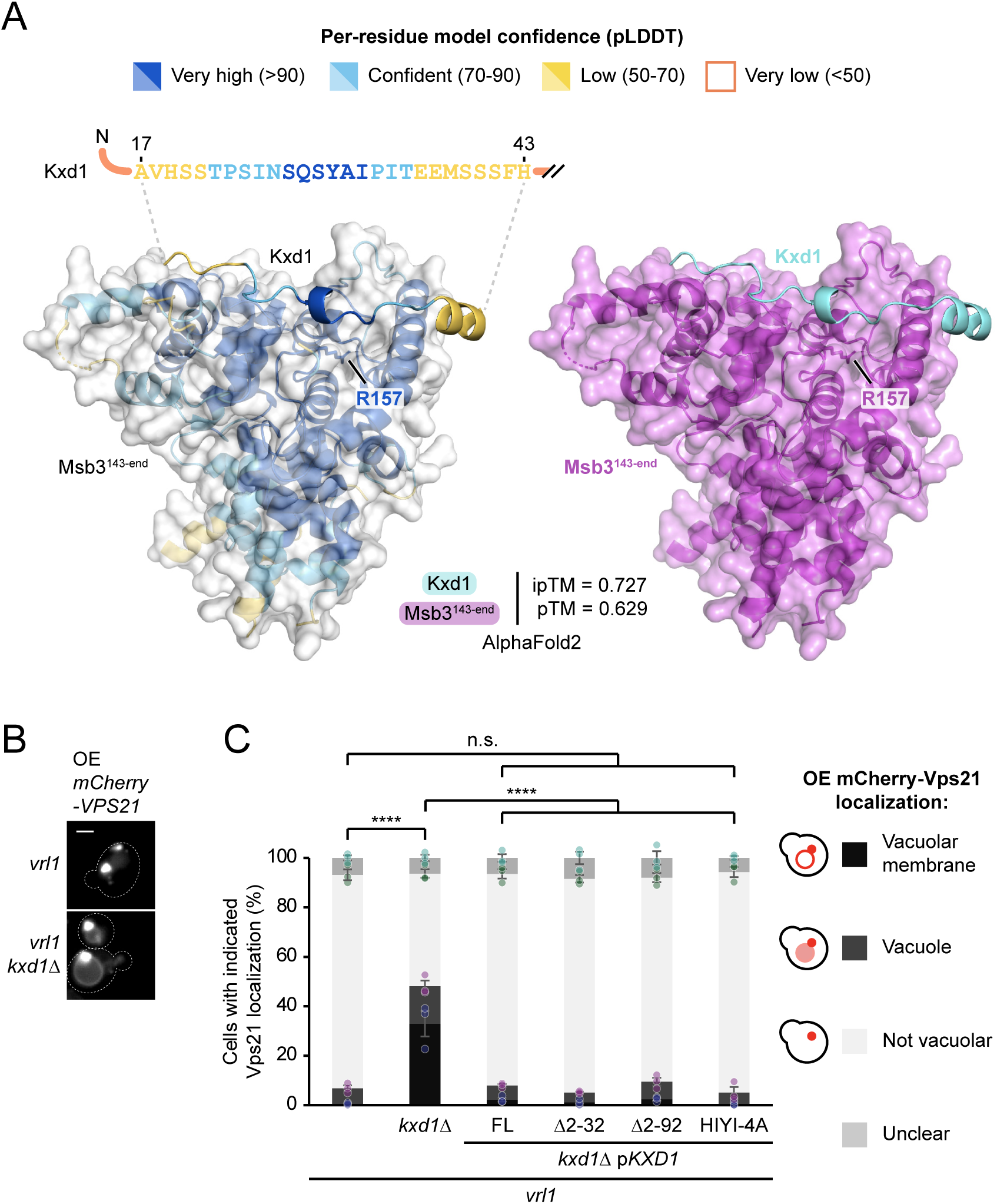
(A) AlphaFold2 confidently predicts an interaction between Msb3 and the unstructured N-terminal domain of Kxd1. AlphaFold2-predicted model of the C-terminal region of Msb3 with full-length Kxd1. Models are coloured by pLDDT (left) and by chain (right); only residues with pLDDT>50 are shown. Sequence of confidently modelled region of Kxd1 is shown coloured by pLDDT. ipTM and pTM scores of predicted structure shown at top right. (B) Deletion of *KXD1* causes Vps21 mislocalization to the vacuolar limiting membrane. Fluorescence micrographs of mCherry-Vps21-overexpressing cells, showing effect of *KXD1* deletion. (C) The predicted Msb3-interacting site in the Kxd1^Nt^ is dispensable for Vps21 inactivation by VINE. Effect of *KXD1* deletion and complementation by plasmid-expressed *KXD1* alleles on localization of RFP-Vps21 in select strains shown in Figure *S5B* and *5H*. Legend shows strategy for manual quantitation of blinded images. One-way ANOVA with Tukey’s multiple comparisons test; n=3, cells/strain/replicate ≥ 153; n.s. = p > 0.05, **** = p < 0.0001. Scale bars, 2 µm. Error bars report SEM. OE, overexpressed. HIYI-4A, H19A I25A Y30A I32A. pLDDT, predicted local-difference distance test.

**Table S1. Vrl1 DHFR Interactors.** Vrl1 WT/mutant ratio Z-scores for top Vrl1 interactors.

**Table S2. List of *Saccharomyces cerevisiae* strains used in this study.**

**Table S3. List of plasmids used in this study.**

**Table S4. List of oligonucleotides used in this study.**

